# Proteostasis control via HSP90α sustains YAP activity to drive aggressive behaviours in cancer-associated fibroblasts

**DOI:** 10.1101/2025.04.04.647210

**Authors:** Silvia Domínguez-García, Javier Rodríguez, Miguel Juliá, Diane Coursier, Carmen Pérez-López, Patricia Carnicero, Ana V. Villar, Alexander von Kriegsheim, Fernando Calvo

**Author notes:** **Author for correspondence**: Fernando Calvo.

## Abstract

Cancers adapt proteostasis to cope with the burden of misfolded proteins, stabilize key signalling nodes and sustain their malignant behaviour. Tumour stroma is subjected to similar stresses, but how they influence its aberrant status remains unclear. We show that tumour stroma presents consistent upregulation of target genes associated to the major misfolding regulator HSP90 in cancer-associated fibroblasts (CAFs), and that HSP90α is required for CAFs to remodel the extracellular matrix (ECM) and promote cancer cell motility and growth. Mechanistically, HSP90α sustains TGFβ responses and YAP protein levels required for CAF functionality. In vivo, stromal or fibroblast-specific loss of HSP90α results in reduced ECM deposition, angiogenesis, growth and dissemination of breast tumours. Clinical analyses reveal a correlation between HSP90-dependent programs and YAP activity in CAFs, that are also associated with poor patient prognosis. Our findings uncover a link between proteostasis, mechanotransduction and generation of aggressive tumour microenvironments through HSP90α.

## MAIN TEXT

Cancer disrupts tissue homeostasis, leading to changes in stromal and immune cells and altering tissue properties. This modified context, defined as the tumour microenvironment (TME), actively contributes to tumour progression, dissemination, and therapy response^1^. Among stromal cells, CAFs are primary modulators of the TME, actively remodelling the ECM, which modulates tissue stiffness and promotes malignant progression^2^**^-^**^4^. Additionally, CAFs generate chemical signals that influence cancer, stromal, and immune cell compartments, promoting tumour growth, invasion, angiogenesis and immune suppression.

To sustain their altered state, malignant cells adapt to both intrinsic and extrinsic stressors by co-opting evolutionarily conserved responses^5^. Many of these involve the adaptation of protein homeostasis (proteostasis), including enhanced chaperone function, to cope with the burden of misfolded proteins, stabilize key signalling nodes, and sustain the hyperproliferative nature of cancers^6^. Tumour stroma and CAFs are usually subjected to similar types of stresses^7^ and recent studies, including our own, have revealed the importance of integrated stress responses^8^, hypoxia responses^9^, and heat shock responses^10, 11^ in this context. Furthermore, CAFs respond to mechanical stress within the TME by activating the mechanotransducer YAP, which promotes ECM remodelling and tissue stiffening^10, 12, 13^.

The heat shock protein 90 (HSP90) chaperone machinery is a key regulator of proteostasis under both physiological and stress conditions in eukaryotic cells^14^. HSP90 assists folding of *de novo* proteins, prevents stress-induced protein misfolding and stabilizes protein levels of multiple client proteins, participating in various processes beyond protein folding. Moreover, HSP90 plays a crucial role in cancer owing to the different types of stresses that exert additional pressure on proteostasis control, and the extensive reliance of cancer cells on HSP90-assisted signalling pathways^15^. Indeed, HSP90 levels are markedly increased in tumours, and HSP90 expression is associated with poor prognosis in breast cancer (BC)^16^. Studies in fibrotic conditions have suggested that HSP90 participates on TGFβ-mediated signalling and fibroblast activation^17, 18^. Nevertheless, the role of HSP90 and proteostasis mechanisms in the adaptive processes of TME components during tumour progression remain elusive.

Here, we demonstrate that stromal reactions in tumours are characterized by the upregulation of HSP90-target genes in CAFs, and that HSP90α depletion impairs CAF-driven ECM remodelling, cancer cell motility, and tumour growth. Mechanistically, HSP90α interacts with YAP and stabilizes its protein expression levels, sustaining mechanotransduction. In vivo, global host and CAF-specific HSP90α deletion is associated with reduced ECM remodelling and angiogenesis, affecting both primary and metastatic tumour development. HSP90α-specific gene signatures correlate significantly with YAP activity and desmoplastic phenotypes in CAFs from human specimens, and stratify BC patients with poorer survival. These findings uncover a novel pro-tumour role of HSP90 through CAF-dependent mechanisms and suggest a proteostatic vulnerability for systems with aberrant YAP activity.

## RESULTS

### HSP90-dependent transcriptional programs are a consistent feature of CAFs associated with aggressive TMEs and poor patient prognosis

In search of novel drug targets for TME normalization, we designed an unbiased chemical genomics approach leveraged on the generation of gene signatures consistently dysregulated in tumour stroma and the interrogation of the Library of Integrated Network-based Cellular Signatures (LINCS) database^19^ (**Fig. 1a**). First, we identified differentially expressed genes (DEG) that were consistently up- or downregulated in tumour stroma from human breast, colorectal and ovarian cancer (**Fig. S1a&b**, **Supplementary Table 1**). This analysis informed of shared gene expression patterns characteristic of tumour stroma, including 209 consistently upregulated DEGs (‘Cancer Stroma Hi signature’)(**Fig. 1b**). We validated that the ‘Cancer Stroma Hi’ signature was also significantly upregulated in independent datasets of BC stroma (**Fig. S1c**). Correlation analysis indicated that ‘Cancer Stroma Hi’ expression in tumour stroma was primarily associated with signatures assigned to aggressive tumours, desmoplastic responses and activated fibroblasts (**Fig. S1d**), suggesting CAFs as the main contributors. CAFs present a level of heterogeneity, with distinct molecular subtypes characterized by the expression of specific gene sets. Using previously described gene signatures for BC CAF subtypes^20^**^-^**^22^, we observed that ‘Cancer Stroma Hi’ expression was mainly associated with the expression of myofibroblast CAFs (myCAF) and desmoplastic CAF subtypes (**Fig. S1f**).

**Figure 1.**
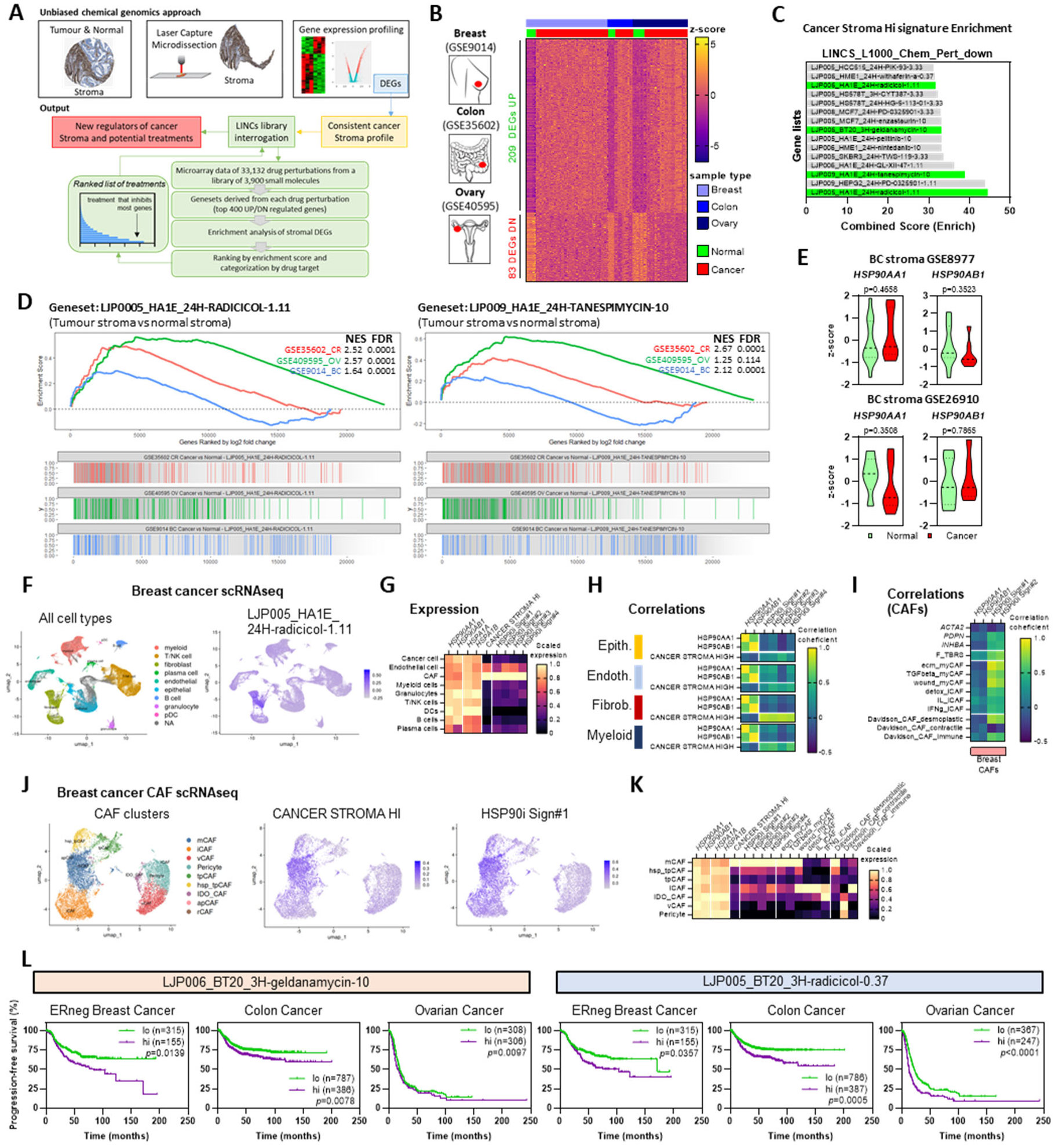
HSP90 signatures are enriched in cancer stroma and primarily associated with CAFs. **A.** Schematic representation of the workflow for the identification of new regulators of cancer stroma with possible therapeutic potential. **B.** Heatmap showing z-scores of consistent upregulated and downregulated DEGs in tumour stroma against normal stroma in breast (GSE9014}, colorectal (GSE3S602} and ovarian (GSE40S9S} patients (Breast: normal, n; 6; cancer, n; 53. Colon: normal, n; **4;** cancer, n; 13. Ovary: normal, n; 8; cancer, n; 31) **C.** Graph showing the signatures with higher combined enrichment score (LINCS_LlOO_Chem_Pert_down database} for the "CANCERSTROMA HI" signature. Signatures related to HSP90i are shown in green. **D.** Graphs showing GSEA plots of HSP90i signatures UPOOOS_HA1E_24H-RADICICOL-l.11 (left} and UP009_HA1E_24H-TANESPIMYCIN (right} in colorectal (GSE35602_CR, red}, ovarian (GSE40595_OV, green} and breast (GSE9014_BR, blue} cancer stroma when compared to normal stroma. Normalized Enriched Score (NES} and False Discovery Rate (FDR} coefficients are also shown. **E.** Graphs showing the z-score value for the gene expression of *HSP90AA1* and *HSP90AB1* in normal and cancer stroma of the BC patient datasets GSE8977 (top panels} and GSE26910 (bottom panels}. (GSE8977: normal, n;lS; cancer, n;7, GSE26910: normal, n:.6; cancer, n:.6}. **F.** UMAP of all cell types identified in BC patients (left} and expression matrix for the UP0OOS_HA1E_24H-RADICICOL-l.11 score in each cell (right} as informed by scRNAseq analysis^23^. Each dot in the graph represents a single cell. **G.** Heatmap showing scaled gene expression values of each HSP90 isoform *(HSP90AA1* and *HSP90AB1),* HSP90 isoform-specific clients *(HSPAlA* and *HSPAlB),* "CANCERSTROMA HI" and all four HSP90i signatures in each cell cluster from (F}. **H.** Heatmap showing correlation coefficients of *HSP90AA1, HSP90AB1* and the "CANCER STROMA HI" signature with all four HSP90i signatures for the indicated cell types from (F). *Epith: epithelial cells. Endoth: endothelial cells. Fibrob: fibroblasts.* **I.** Heatmap showing correlation coefficients of *HSP90AA1, HSP90AB1,* and HSP90i signatures Ill and #2 with indicated genes *(ACTA2, PDPN* and *INHBA)* and different CAF-related signatures in CAFs from BC scRNAseq dataset^23^. **J.** UMAP of CAF clusters identified in BC patients (left) and expression matrices for the "CANCER STROMA HI" (middle) and UPOO0S_HA1E_24H-RADICICOL-l.11 (’HSP90i Sign#l’, right) signatures in each individual cell, as informed by scRNAseq analysis. Each dot in the graph represents a single cell. **K.** Heatmap showing scaled gene expression values of indicated genes and gene signatures in the different CAF clusters from BC patients from (J). **L.** Graphs showing the percentage of progression-free survival in patients with high (hi) or low (lo) expression of HSP90i signatures LP006_BT20_3H-geldanamycin-10 (left) and UPOOS_BT20_3H-radiciol-0.37 (right), for the ER negative breast, colon and ovarian cancer patients. The p-value for each dataset and number of patients in each group is included in the corresponding graph. *HSP90i Sign#l: LJP005_HA1E_24H-radicicol-1.11. HSP90i Sign#2: LJP009_HA1E_24H­ tanespimycin-10. HSP90i Sign#3: LJP006_BT20_3H-geldanamycin-10. HSP90i Sign#4: LJP006_HA1E_24H-radicicol-l.11*.

Next, we queried the ‘Cancer Stroma Hi’ signature against the LINCS database, focusing in gene sets for perturbations that result in gene expression inhibition (‘LINCS_L1000_Chem_Pert_down’). Several HSP90 inhibitors (HSP90i) were found among the top-ranked small molecules whose gene-expression profiles were correlated to the ‘Cancer Stroma Hi’ profile (**Fig. 1c**). These include the natural antibiotics Radicicol and Geldamycin, as well as the derivate product Tanespimycin (also known as 17-AAG). In agreement, gene set enrichment analysis (GSEA) showed a significant enrichment of HSP90i signatures in cancer stroma from breast, colon and ovarian cancer (**Fig. 1d**). In mammalian cells, HSP90 functions are orchestrated by two major cytoplasmic isoforms: HSP90α (encoded by *HSP90AA1*) and HSP90β (encoded by *HSP90AB1*). Of note, we did not observe significant differences in the gene expression of both isoforms between normal and tumour stroma (**Fig. 1e**). All four HSP90i signatures were correlated with signatures of aggressive tumours, desmoplastic responses, activated fibroblasts and myCAFs (**Fig. S1h&i**). These observations were further validated using independent scRNAseq data from BC patients^23^ that showed: (i) preferential expression of HSP90i signatures in CAFs (**Fig. 1f&g**); (ii) high correlation between ‘Cancer Stroma Hi’ and HSP90i signatures in CAFs (**Fig. 1h**); and (iii) high correlation between HSP90i signatures and myCAF characteristics (**Fig. 1i**). Examining the distribution of signatures in the UMAP projection of CAF populations from the BC scRNAseq^23^ (**Fig 1j**) showed that ‘Cancer Stroma Hi’ and HSP90i signatures were expressed at elevated levels in the mCAF/myCAF subtype (**Fig 1j&k**). Of note, clusters characterised by high expression of heat-shock proteins and termed ’heat-shock protein-high tumour CAFś (hsp_tCAFs)^23^, had lower levels of ‘Cancer Stroma Hi’ and HSP90i signatures, emphasizing the divergence between HSP90 gene expression and HSP90-dependent transcriptional programs. Furthermore, we observed a significant lower progression-free survival in breast (ER-negative), colon and ovarian cancer patients that present higher levels of HSP90i signature expression (**Fig. 1l**). These results indicate that the HSP90 target genes found consistently upregulated in the tumour stroma have clinical value and correlate with poorer patient prognosis.

To validate the relevance of HSP90 activity in modulating CAF phenotypes, we turned to tractable models of normal fibroblasts (NFs) and CAFs (**Fig. S2a**). GSEA of CAF vs NFs from different systems (including our model of murine BC CAF/NFs) indicated a consistent and significant upregulation of HSP90i signatures in CAFs (**Fig. S2b**). Since transcriptional programs may be context dependent and LINCS HSP90i signatures were obtained mainly from cancer models, we queried an HSP90i signature obtained in mouse embryonic fibroblasts (MEFs)^24^ (**Fig. S2a**). Notably, this fibroblast-specific HSP90i signature was also significantly enriched in CAFs compared to NFs (**Fig. S2c**). Similar findings were observed when we queried a gene expression dataset of freshly isolated murine CAFs compared to

NFs (**Fig. S2d**), thus ruling out in vitro artefacts contributing to these characteristics. Murine and human BC CAFs had no significant differences in the expression of *HSP90AA1*, *HSP90AB1* or genes associated to HSP90 function compared to NFs (**Fig. S2e&f**). Interestingly, Mass Spectrometry (MS) proteomic analysis of our murine model of BC CAFs and NFs^12^ informed that the protein content of Hsp90α is slightly higher in CAFs, which is compensated with a decrease in Hsp90β expression, with no overall differences in total Hsp90 expression (**Fig. S2g**). No differences were observed after TGFβ1 treatment of NFs, despite the documented role of HSP90 in modulating TGFβ signalling^25^. Altogether, these findings suggest that CAFs present gene expression programs highly dependent on HSP90 function.

### *Hsp90aa1*-depleted CAFs display reduced pro-tumoral activities

To assess the role of HSP90 in modulating CAF behaviour, we designed a battery of functional assays employing two stablished systems: murine CAFs that have a persistent activated status and mostly associated to myCAF phenotypes^12^; and murine NFs stimulated with TGFβ1 to promote myofibroblast activation (**Fig. S3a**). Although both HSP90 isoforms bind equally to their cochaperones and perform similar molecular functions, they have differential effects on substrate interactions that may impact in distinct molecular and cellular processes^14, 25^. To evaluate which isoform affected CAF functionality, the aforementioned models were subjected to RNAi mediated gene silencing for both *Hsp90aa1* (Hps90α) and *Hsp90ab1* (Hsp90β). In addition, we isolated normal mammary gland fibroblasts from *Hsp90aa1* knock-out mice (NF^KO^) and wild-type littermates (NF^WT^). We confirmed the efficacy and specificity of silencing at the level of gene (**Fig. 2a and S3b&c**) and protein expression (**Fig. S3d&f**) in all systems.

**Figure 2.**
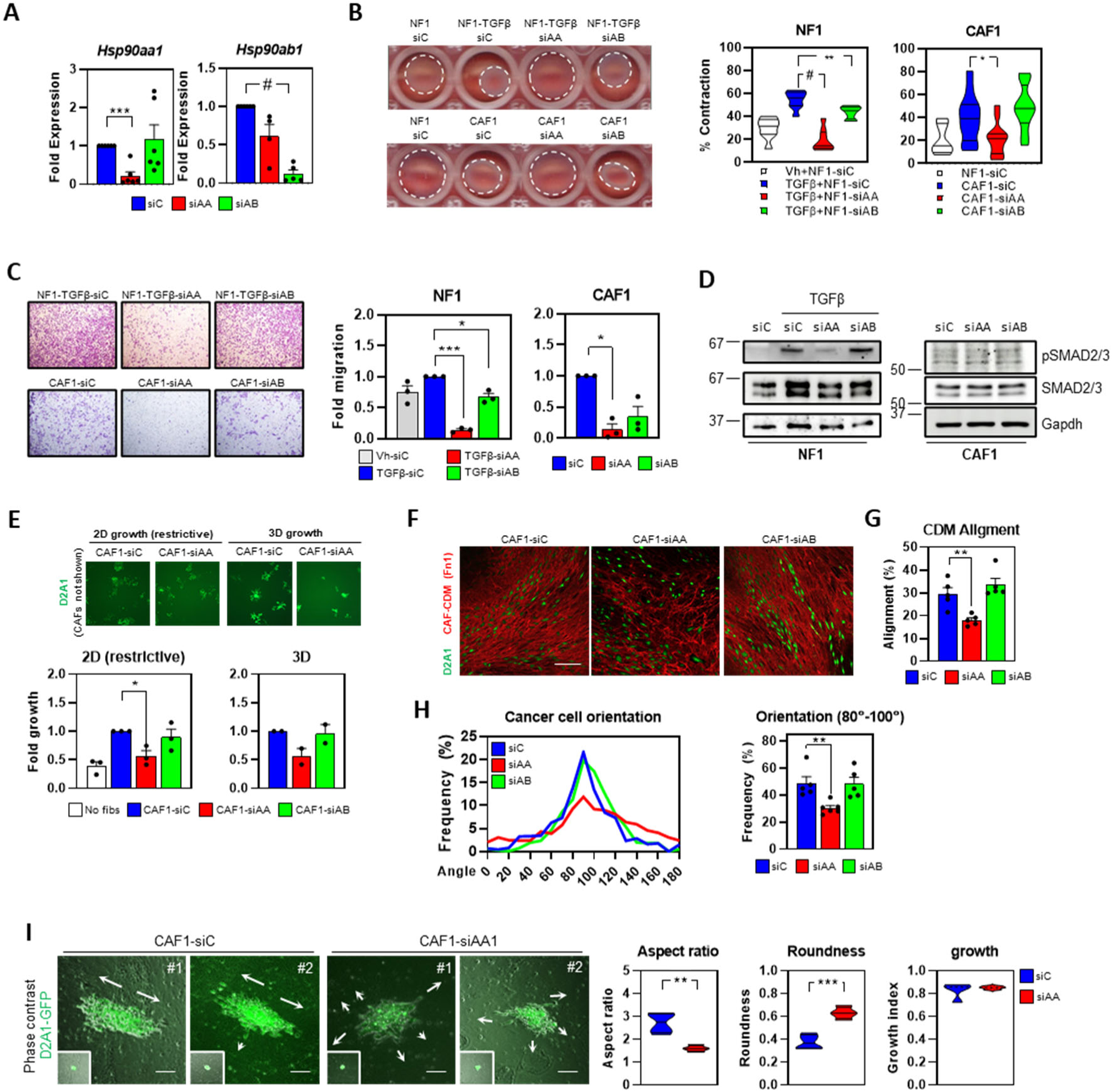
HspSOaai-depleted CAFs display reduced pro-tumoral activities. **A.** Graphs show *Hsp90oal* and *Hsp90obl* fold mRNA expression (relative to *Gapdh)* in murine CAF1 after transfection with control (siC), *Hsp90aal* (siAA) or *Hsp90abl* (siAB) RNAi (smart-*poo\s)(Hsp90aal:* all, n=6. *Hsp90abl:* sic, n=6; siAAl, n=4; siABl, n=5). **B.** Images show representative gel contraction assays for murine NF1 and CAF1 after transfection with control (siC), *Hsp90aal* (siAA) or *Hsp90abl* (siAB) RNAi (smart-pools). Where indicated, NF1 were stimulated with vehicle (Vh) or TGFgl. Graphs show percentage of gel contraction in the indicated conditions (NF1: all, n=8. CAF1: NFl-siC, n=8; rest, n=12). **C.** Images show representative crystal violet-stained cells that migrated in transwell assays for murine TGFfSl-stimulated NF1, and unstimulated CAF1 after transfection with control (siC), *Hsp90aal* (siAA) or *Hsp90abl* (siAB) RNAi (smart­pools). Graphs show fold migration for the indicated conditions, including unstimulated NFl-siC (n=3). **D.** Representative Western blots showing expression levels of phospho-SMAD2/3, total SMAD2/3 and Gapdh in murine NF1 and CAF1 after transfection with control (siC), *Hsp90aal* (siAA) or *Hsp90abl* (siAB) RNAi (smart-pools). Where indicated, NF1 were stimulated with TGFpi. **E.** Images show D2A1-GFP cells (green) co-cultured with CAF1 (not shown) after transfection with control (siC), *Hsp90aal* (siAA) or *Hsp90abl* (siAB) RNAi (smart-pools). Images on the left show results in 2D restrictive conditions; images on the right show results on top of matrigel layers (3D conditions). Graphs show corresponding fold growth of D2A1 for the indicated conditions, including no fibroblast control. (2D: n=3. 3D: n=2). **F.** Representative images of D2A1-GFP (green) cultured in CDMs generated by murine CAF1 after transfection with control (siC), *Hsp90aal* (siAA) or *Hsp90abl* (siAB) RNAi (smart-pools). CDM architecture was detected by Fnl staining (red). Scale bar, 150 pm. G. Graph shows alignment score of the Fnl fibres of indicated conditions from (F). (n=5). **H.** *(Left)* Graph showing the distribution frequency for the different orientation angles in which the D2A1-GFP were disposed on top of the indicated CAFl-derived CDMs. The corresponding angles were determined using the frequency peak as reference (9(5). *(Right)* Histogram showing percentage of D2A1-GFP cells oriented between 8(5-100° angles (n=5). **I.** Images show two different D2A1-GFP spheroids (green) at the final point (main image) and starting point (small insert) when incubated over CDMs (phase contrast image on greyscale) from murine CAF1 after transfection with control (siC) or *Hsp90aal* (siAA). White arrows indicate the preferred direction of cancer cell migration. Graphs show the aspect ratio, roundness and growth index of D2A1 spheroids at the final point for each condition (n=4). Scale bar, 150 pm. Where indicated, individual *p* values are shown; alternatively, the following symbols were used to describe statistical significance: *\*,P<* 0.05; **, P < 0.01; ***, P < 0.001; *it, P* < 0.0001.

Analysis of gel contraction assays, a measure of the contractile ability of CAFs associated with ECM remodelling and cancer cell invasion^12^, showed that silencing of *Hsp90aa1* reduced this function in both murine CAF and TGFβ1-stimulated NFs (**Fig. 2b and Fig. S3f**), as well as in NF^KO^ compared to NF^WT^ (**Fig. S3g**). *Hsp90aa1* silencing also negatively affected the migratory capacities of CAF1 and TGFβ1-stimulated NF1, with minor differences after *Hsp90ab1* depletion in TGFβ1-stimulated NF1 (**Fig. 2c**). Unstimulated NF^KO^ also presented less migration when compared to NF^WT^ (**Fig. S3h**). These results suggested a specific role for HSP90α in modulating fibroblast behaviour that was not shared with HSP90β. It has been shown that HSP90α potentiates TGFβ signalling by stabilizing TGFβ receptors in non-tumoral contexts^18, 25^. Given the relevance of this pathway in CAF emergence, we explored the effect of *Hsp90aa1* silencing in TGFβ/SMAD signalling in our models. We observed that depletion of *Hsp90aa1* but not *Hsp90ab1* reduced SMAD2/3 activation in TGFβ1-stimulated NF1 (**Fig. 2d**). Interestingly, we did not observe such effect in unstimulated CAFs, which were also unaffected by TGFβ inhibitor treatment on their ability to contract collagen gels (**Fig S3i**). Moreover, we did not observe cell cycle deficiencies after *Hsp90aa1* or *Hsp90ab1* silencing (**Fig. S3j**), ruling out potential effects on cell viability. These results suggest that HSP90α is not operating via TGFβ signalling in CAFs, and that additional mechanisms might be at play. Focusing on CAFs, we also observed that knocking-down *Hsp90aa1* reduced the ability of CAFs to promote growth of murine BC models in restrictive (*i.e.* low nutrient availability) and in 3D conditions (**Fig. 2e**), which was also observed in the knock-out system (**Fig. S3k**). We next proceeded to evaluate the role of HSP90 in modulating functions associated with CAF/myCAFs such as ECM deposition employing cell-derived matrices (CDMs)(**Fig. S4a**). These analyses informed that *Hsp90aa1* silencing affected the ability of CAFs to generate isotropic CDMs, reducing the total length and overall alignment of the ECM fibres (**Fig. 2f&g and Fig. S4b&c**). Isotropic ECMs have been associated with increased cancer cell invasion, as cancer cells employ the aligned ECM fibres for fast orientated migration^26, 27^. In agreement, D2A1 murine BC cells growing on CDMs generated by *Hsp90aa1*-depleted CAFs presented more heterogeneous patterns, whereas cells growing on control and *Hsp90ab1*-silenced CAF CDMs were more orientated (**Fig. 2f&h**). Similar findings were observed with alternative CAF lines (**Fig. S4d**). Moreover, when cancer cell spheroids were seeded over control CDMs they grew and migrated on a preferential direction, whereas spheroids over *Hsp90aa1*-depleted CDMs migrated in a disorganized fashion (**Fig. 2i and Supplementary Movies 1-4**).

To assess if HSP90α was affecting the activated status of CAFs, we investigated the expression levels of well-characterized markers. This analysis showed that many genes showed no relevant differences after *Hsp90aa1* or *Hsp90ab1* modulation in CAF1 and TGFβ1-stimulated NF1 (**Fig. S4e-g**). There were few exceptions including: *Pdpn*, *Inhba* (CAF markers); *Has2* (ECM deposition/remodelling); and *Rock1* and *Sept9* (cytoskeleton), all of which were specifically downregulated by *Hsp90aa1* RNAi. In addition, we detected several gene expression responses associated to *Hsp90aa1* depletion specific for TGFβ1-stimulated NFs and not CAFs (*Tgfb1*, *Fgf2*, *Pdgfa* and *Pdgfb*).

Taken together, our data confirms that HSP90α loss results in reduced sensitivity to TGFβ stimulation in fibroblasts, as observed in other non-tumoral contexts^18, 25^. In addition, our results suggest that *Hsp90aa1* depletion in activated fibroblasts impairs signalling cascades required for their pro-tumour behaviour in a TGFβ-independent manner.

### *Hsp90aa1* depletion in activated fibroblasts inhibits YAP-dependent transcriptional programs

To shed light into the mechanisms employed by HSP90 to modulate pro-tumour behaviours in CAFs, we performed transcriptomic and MS proteomic analyses of CAFs and TGFβ1-stimulated NFs after transfection with control, *Hsp90aa1* and *Hsp90ab1* RNAi. Principal Component analysis (PCA) of RNAseq data showed that *Hsp90aa1*-depleted CAFs presented a distinct transcriptional landscape apart from control and *Hsp90ab1*-silenced CAFs (**Fig. 3a**), in line with functional observations. This was also evidenced by the number of up- or downregulated DEGs in *Hsp90aa1*- and *Hsp90ab1*-silenced CAFs compared to control (**Fig. 3b and Fig. S5a**). Enrichment analysis of DEGs showed no relevant associations for common genes (**Fig. S5b**). However, DEGs specifically downregulated in *Hsp90aa1*-silenced CAFs showed enrichment in gene sets associated with cell cycle, DNA repair and proteasome regulation (**Fig. S5b&c**). Interestingly, when we queried *Hsp90aa1* DEGs against gene sets associated with processes related to fibroblast activation/CAFs, we observed a high level of enrichment for signatures related to activation of YAP/TAZ in CAFs (‘CAF_YAPTAZ_ALL_UP’), at relatively similar levels to signatures associated to HSP90i responses in MEFs (‘MEF-HSP90i signature’)(**Fig. S5d**). This prompted us to perform GSEA, which showed that *Hsp90aa1* depletion in CAFs significantly downregulates gene signatures associated to YAP/TAZ and RhoA activity, desmoplastic CAFs (‘DAVIDSON_CAF-S2_DESMOPLASTIC_UP’) and Src activities (‘DASATINIB_DN’)(**Fig. 3c&d and Fig. S5e**). Since YAP/TAZ activity can be associated with proliferative transcriptional responses in certain systems^28^, it was plausible that the emergence of this signature was a consequence of cell cycle defects, albeit these were also significantly downregulated by *Hsp90ab1* depletion and we did not observe major changes in cell cycle (**Fig. S3i**). Nevertheless, we further confirmed the dysregulation of YAP/TAZ signatures in *Hsp90aa1*-depleted CAFs by querying our YAP/TAZ signature devoid of E2F genes (‘NON_PROLIF_CAF_ALL_UP’) (**Fig. 3c&d and Fig. S5e**). Noteworthy, TGFβ signatures presented no significant differences in *Hsp90aa1-* or *Hsp90ab1-depleted* CAFs. Interestingly, we obtained similar results when we analysed the RNAseq dataset of TGFβ1-stimulated NFs (**Fig. 3d and Fig. S6a-g**). Of note, in the TGFβ1-stimulated NF system, HSP90α but not HSP90β modulation resulted in the downregulation of TGFβ-dependent signatures, in line with our previous results (**Fig. 2d**).

**Figure 3.**
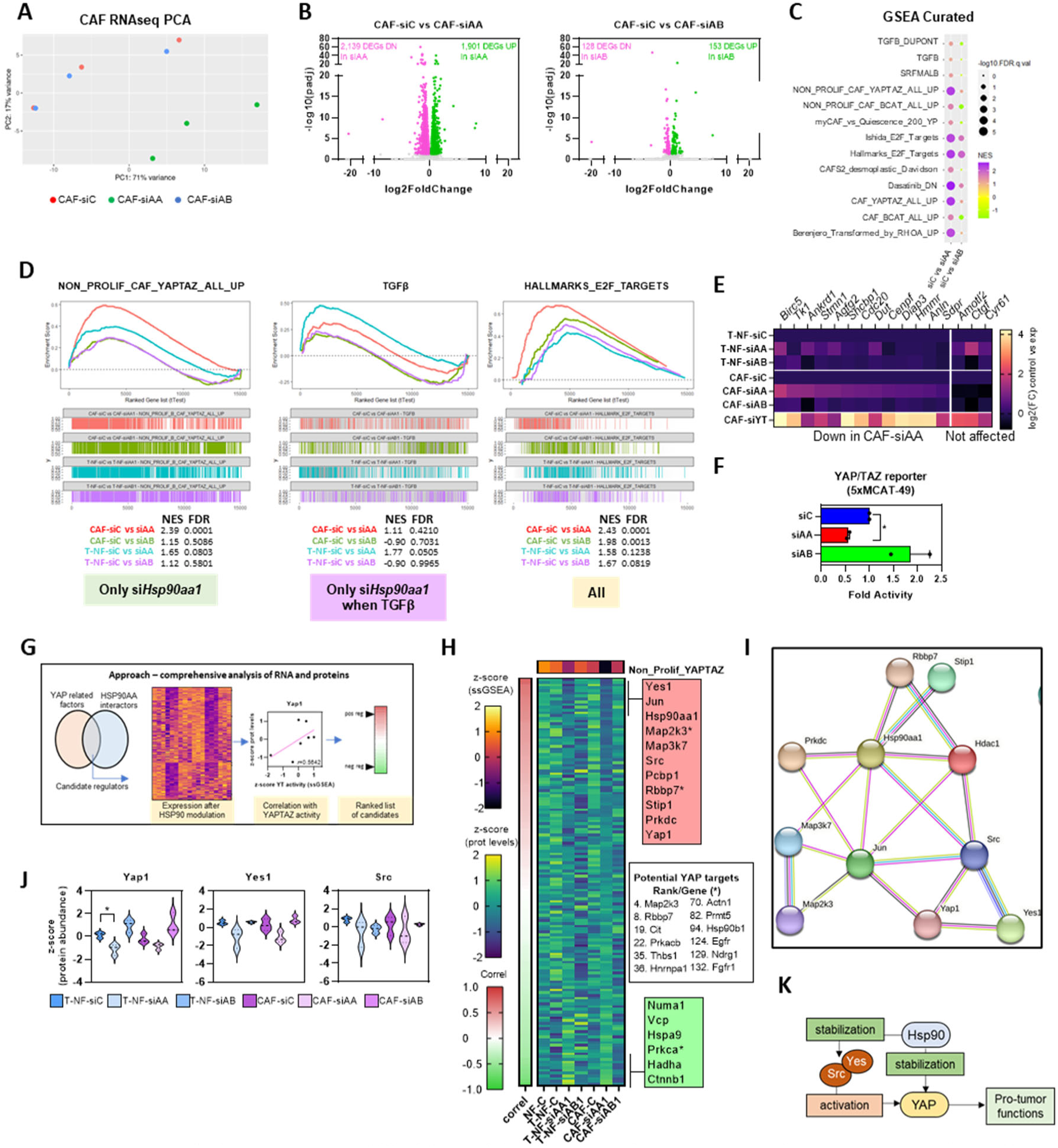
*Hsp90aal* depletion in activated fibroblasts inhibits YAP-dependent transcriptional programs. **A.** Graph showing PCA based on RNAseq of murine CAFl after transfection with control (siC), *Hsp90aal* (siAA) or *Hsp90abl* (siAB) RNAi (smart-pools). Each dot represents an independent replicate (n=3). **B.** Graphs showing the upregulated (green dots) and downregulated (pink dots) DEGs as determined by their -LoglO P adjusted value (Padj) and the Log2 fold change, when comparing *Hsp90aal-depleted* (left) and *Hsp90abl-depleted* (right) CAFl cells, in comparison with control (siC). Each dot represents an individual gene. **C.** Bubble plot summarizing GSEA results of "curated gene sets" when comparing CAFl control (siC) vs *Hsp90aa1-depleted* (siAA) or *Hsp90ab1-* depleted (siAB) CAFl. The size of the bubbles represents the level of significance of the signature (-Logl0 FDR), and their colour represents the level of NES. **D.** GSEA plots show the level of activity of the indicated signatures in *Hsp90aal-depleted* CAFl compared with CAFl control (red line), *Hsp90abl-depleted* CAFl compared with the CAFl control (green line), *Hsp90aal-depleted* TGFSl­ stimulated NFl compared with the TGFSl-stimulated control (blue line) and *Hsp90abl-depleted* TGFSl-stimulated NFl compared with the TGFSl-stimulated control (purple line). NES and FDR coefficients are also indicated. Where indicated, 5 ng/ml of TGFSl was added for 24 h. **E.** Heatmap showing the log2(FC) values of expression of indicated YAP target genes when comparing control settings against indicated RNAi perturbations. On top, values for murine TGFSl-stimulated NFl cells after transfection with control (siC), *Hsp90aal* (siAA) or *Hsp90abl* (siAB) RNAi (smart-pools). On the bottom, values for murine CAFl cells after transfection with control (siC), *Hsp90aa1* (siAA), *Hsp90ab1* (siAB) or *Yapl* and *Taz* (siYT) RNAi (smart-pools) (n=3). Data from siYTwas obtained from^10^. **F.** Graph shows luciferase activity of YAP/TAZ activation (SxMCAT-49-lux reporter) in CAFl after transfection with control (siC), *Hsp90aa1* (siAA) or *Hsp90ab1* (siAB) RNAi (smart-pools). (n=2). **G.** Schematic representation of the workflow followed for the integration of the transcriptomic and proteomic datasets. **H.** Heatmap showing mean z-score protein expression of candidate factors selected as described in (F) for murine NFl and CAFl after transfection with control (siC), *Hsp90aal* (siAA) or *Hsp90ab1* (siAB) RNAi; where indicated (T), NFl were stimulated with TGFSl. Factors are ranked based on their correlation values with the "NON_PROLIF _ YAPTAZ" signature (ssGSEA values, z-score normalized, top heatmap). The top positive correlated factors are indicated in a green panel, whereas the top negative correlated factors are presented in a red panel. Some of the known YAP targets have been noted in the white right panel with their corresponding position in the rank (n=3 for both protein and ssGSEA, all conditions). I. Diagram showing the interaction grid containing the top positive correlated proteins from the ranked list in (H). **J.** Graphs show z-score normalized protein expression of Yapl, Yesl and Src for the indicated conditions in (H)(n=3). **K.** Schematic representation of the proposed mechanism of YAP regulation by HSP90a, based on analyses summarized in (G). Where indicated, individual *p* values are shown; alternatively, the following symbols were used to describe statistical significance: •, *P* < 0.05.

YAP/TAZ are transcriptional regulators with paramount roles in development and cancer^29^. In CAFs, YAP/TAZ are activated in response to mechanical cues and, in turn, YAP establishes a transcriptional program that enhances CAF functionality ^10, 12, 13^. Thus, our results suggested that HSP90α may operate via YAP to regulate CAF behaviour. We confirmed that several YAP/TAZ target genes in CAFs were downregulated after *Hsp90aa1* knock-down including *Diap3* and *Anln* (**Fig. 3e**), which we previously showed to play relevant roles in CAF functions^12^. This was concomitant to a reduction in YAP transcriptional activity after *Hsp90aa1* silencing in CAFs (**Fig. 3f**). Given the role of HSP90 in modulating protein folding and stability, we performed proteomics analyses to explore how HSP90 may be affecting YAP/TAZ activity. In line with our transcriptomic results, *Hsp90aa1* silencing in both CAFs and TGFβ1-stimulated NFs affected YAP/TAZ targets such as Diaph3, Anln and Birc5 (**Fig. S7a&b**). In agreement, GSEA applied to proteomic data informed of the inhibition of YAP/TAZ and associated mechanotransduction pathways upon *Hsp90aa1* depletion in CAFs and TGFβ1-stimulated NFs (**Fig. S7c**). In the case of TGFβ signatures, these were downregulated by both RNAi but only in the TGFβ1-stimulated NF system.

YAP/TAZ are the main downstream effectors of the Hippo pathway, with critical roles in cell growth and differentiation^30^. The mammalian Hippo pathway consists of a kinase cascade in which MST1/MST2 or MAP4K kinases phosphorylate LATS1/2 (**Fig. S7d**). In turn, activated LATS inhibits

YAP/TAZ through direct serine phosphorylation, preventing nuclear localization and transcriptional activity^31^. Noteworthy, heat-shock stresses can inhibit LATS kinases by promoting HSP90-dependent inactivation of LATS, which potentiates YAP activity^32^. To explore the role of Hippo components in HSP90α-dependent YAP/TAZ modulation, we assessed their protein levels using the proteomics dataset. This analysis showed no relevant or consistent differences between control and *Hsp90aa1*-depleted conditions (**Fig. S7e&f**). This is in line with previous reports showing Hippo-independent regulation of YAP/TAZ in CAFs^10, 12^. In addition, we observed that *Hsp90aa1* knock-down still had inhibitory effects on gel contraction assays on NFs stably expressing a Hippo-insensitive constitutively active YAP mutant (YAP^MUT^)(**Fig. S7g**). Taken together, these results suggest a Hippo-independent mechanism implicated in HSP90α-dependent YAP/TAZ regulation in CAFs.

To identify candidate factors implicated in this process, we designed an approach integrating our transcriptomic and proteomics datasets to identify proteins affected by HSP90α modulation that correlated with YAP/TAZ activity (**Fig. 3g**, *see Methods for details*). This analysis identified some potential YAP targets as top hits (Map2k3, Rbbp7) as well as a core regulatory network dominated by HSP90α and involving Yap1 and the tyrosine kinases Src and Yes1, which can phosphorylate YAP and potentiate its transcriptional activity^33^ (**Fig. 3h-j**). Interestingly, our analysis did not detect TAZ as a potential target of HSP90α, despite the high similarities with YAP. We noticed that TAZ was not among the candidates that presented evidence of HSP90α interaction and therefore was not included in the initial analysis. We assessed the protein levels of Taz in our system and observed a high level of correlation with TAZ-specific (not YAP) gene signatures (**Fig. S7&i**). Noteworthy, TAZ is also subjected to positive upstream regulation by Yes1 and Src^34, 35^, although it has limited functional relevance in CAFs^12^. These results suggest that HSP90α may be modulating CAF behaviour by regulating YAP activity directly, promoting their stabilization, as well as indirectly promoting its activation through tyrosine phosphorylation via Yes1/Src (**Fig. 3k**).

### HSP90α stabilizes YAP protein levels

Activated fibroblasts and CAFs present aberrant activation of signalling cascades that promote pro-tumour behaviours, such as mechanotransduction pathways involving YAP/TAZ and Src^12^. Based on our results, we speculated that YAP/TAZ are client proteins of HSP90α that rely on its function to maintain their high levels of activity, making CAFs particularly sensitive to HSP90α silencing (**Fig. 4a**). To test this possibility, we confirmed that *Yap1* and *Taz/Wwtr1* gene expression levels were not changed by *Hsp90aa1* or *Hsp90ab1* knock-down in CAFs (**Fig. 4b**). However, protein levels of Yap/Taz were reduced after *Hsp90aa1* silencing in CAFs (**Fig. 4c**), as well as in TGFβ1-stimulated NFs (**Fig. S8a&b**). Similar findings were observed in the NF^WT^/NF^KO^ system (**Fig. S8c**). Relative Hippo pathway-induced S127 phosphorylated levels of YAP were not affected by neither treatment (**Fig. S8d**), supporting a Hippo-independent role of HSP90 in CAFs. We and others have described that YAP stability in CAFs is also reinforced by canonical Wnt signalling, which also stabilizes β-catenin^10, 36^. To rule out a role of HSP90α in this regulatory mechanism, we investigated β-catenin levels and observe no differences after *Hsp90aa1* silencing in CAFs (**Fig. S8e**). Furthermore, pulse-chase studies with cycloheximide (CHX) confirmed that YAP and TAZ protein stability was affected in the absence of HSP90α in CAFs (**Fig. 4d**), in TGFβ1-stimulated NFs (**Fig. S8f**) and in the NF^WT^/NF^KO^ system (**Fig. S8g**). Of note, reduced protein stability was also observed in constitutively expressed YAP^MUT^ in the absence of *Hsp90aa1* (**Fig. S8h**). These results confirmed that HSP90 regulation of YAP is not at transcriptional level, and that is independent of Hippo pathway regulation. YAP protein levels were also sensible to HSP90i (Tanespimycin and Geldanamycin) treatment, supporting a role of YAP as a HSP90 client (**Fig. 4e**). Lastly, we confirmed that endogenous YAP was co-immunoprecipitating with endogenous HSP90α. In a first approach, we employed genetically engineered NF and CAF cell lines that have incorporated a V5-dTAG sequence in frame at the C-terminus of the *Yap1* locus (*See Methods for details*). The product of this modified version can therefore by detected and immunoprecipitated with V5 antibodies, and degraded with dTAG (**Fig. S8i**). Employing these tools, we detected that Hsp90 was co-immunoprecipitating with Yap1-V5 (**Fig. 4f and Fig. S8j**). Noteworthy, we observed that co-immunoprecipitated Hsp90 required the expression of Yap1-V5-dTAG, that was more abundant after TGFβ1 stimulation in both NF and CAFs, and that was higher in CAFs than in NFs (**Fig. S8k**). Conversely, immunoprecipitating HSP90 in NF^WT^ enabled the detection of both YAP and TAZ bands, that were completely absent in NF^KO^ cells (**Fig. 4g**).

**Figure 4.**
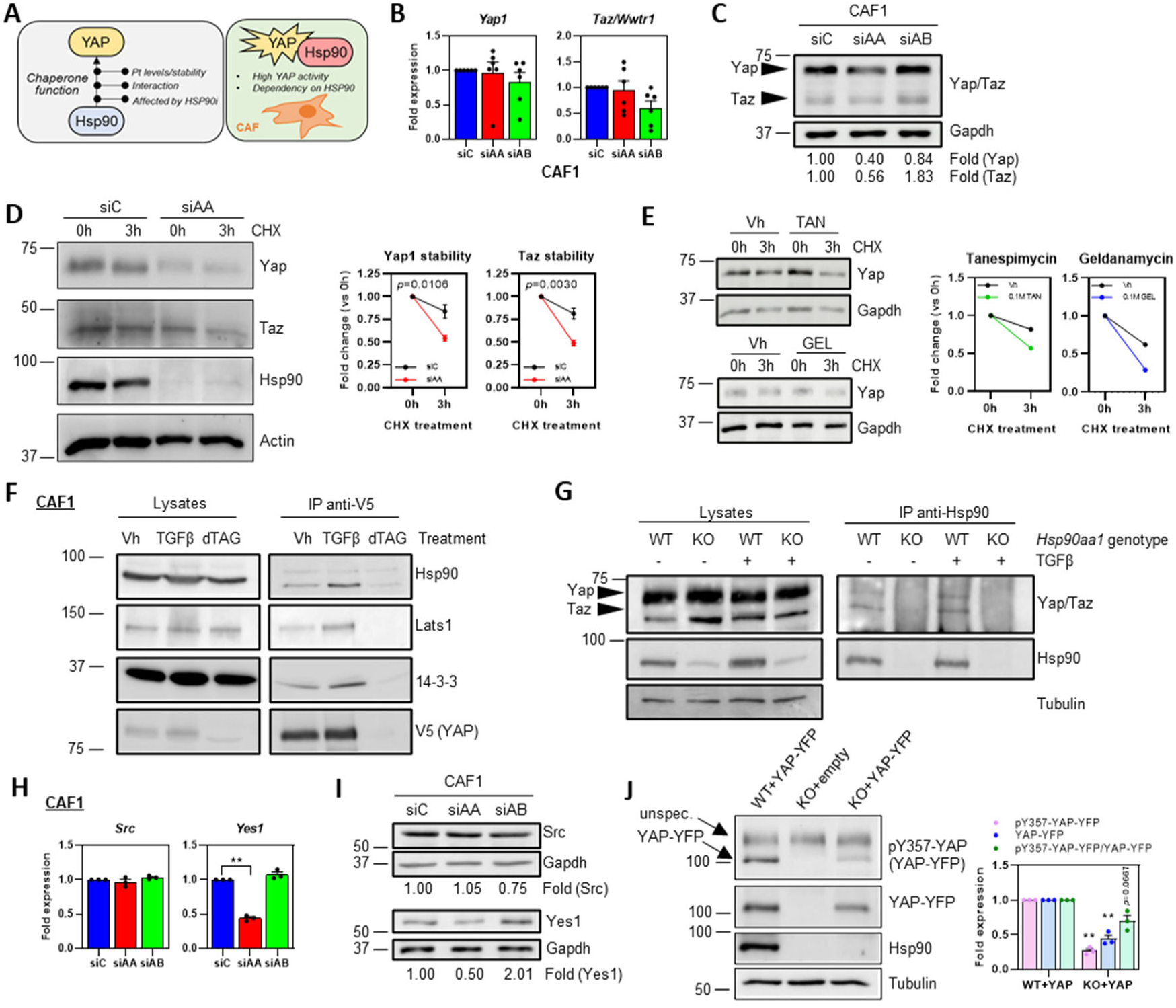
HSP90ct stabilizes YAP protein levels. **A.** Diagram representing the potential regulatory mechanisms linking HSP90 and YAP activities in CAFs. **B.** Graphs showing fold change gene expression of *Yapl* and *Taz/Wwtrl* (relative to *Gapdh)* in murine CAF1 after transfection with control (sic), *Hsp90aal* (siAA) or *Hsp90abl* (siAB) RNAi (smart-pools) by qPCR (n=6). **C.** Western blot showing expression of Yap, Taz and Gapdh in murine CAF1 after transfection with control (siC), *Hsp90aal* (siAA) or *Hsp90abl* (siAB) RNAi (smart-pools). Fold expression levels (relative to Gapdh) are shown. **D.** Western blot showing Yap, Taz, Hsp90 and Actin in murine CAF1 after transfection with control (siC) or *Hsp90aal* (siAA) RNAi (smart-pools) during pulse-chase with CHX at 0 h and 3 h. Graphs represents quantification of the indicated blots (relative to Actin) at 3 h of CHX treatment relative to the expression at 0 h for both conditions (n=4). **E.** Western blots showing Yap and Gapdh expression in CAF1 pre-treated with HSP90i Tanespimycin (TAN) or Geldanamycin (GEL) and subjected to pulse-chase experiments with CHX at 0 h and 3 h. Graphs show quantification of Yap expression (relative to Gapdh) at 3 h of CHX treatment relative to the expression at 0 h for both conditions. **F.** Western blots show co­immunoprecipitation of Hsp90, Latsl and 14-3-3 with V5-YAP (anti-V5) in CAFl-V5-Yap-dTAG cells treated with vehicle (Vh), TGFgl or dTAG. Corresponding total lysates for all antibodies are also shown. **G.** Western blots show co-immunoprecipitation of Yap and Taz with Hsp90 in NF^and NF^K0^ cells. Where indicated, cells were subjected to TGFfl stimulation for 24 h. Corresponding total lysates for all antibodies and Tubulin are also shown. **H.** Graphs show *Src* and *Yesl* fold mRNA expression in murine CAF1 after transfection with control (siC), *Hsp90aal* (siAA) or *Hsp90abl* (siAB) RNAi (smart-pools). Data from RNAseq (n=3). **I.** Western blots show expression levels of Src, Yesl and Gapdh in murine CAF1 after transfection with control (siC), *Hsp90aal* (siAA) or *Hsp90abl* (siAB) RNAi (smart­pools). Fold expression levels (relative to Gapdh) are indicated. J. Western blots show expression of YAP-YFP, tyrosine phosphorylated YAP-YFP (Y357), Hsp90 and Tubulin in NF^WT^ and NF^K0^ cells stably expressing a human YAP-YFP construct, and NF^K0^ expressing an empty vector. Bands corresponding to YAP-YFP and an unspecific band ("unspec.") for the pY357-YAP immunoblot are indicated. Graph shows the fold expression levels of pY357-YAP-YFP and YAP-YFP (relative to tubulin), and pY357-YAP-YFP (relative to YAP-YFP) in NF^WT^-YAP-YFP and NF^K0^-YAP-YFP cells (n=3). Where indicated, individual *p* values are shown; alternatively, the following symbols were used to describe statistical significance: **, P<0.01.

Our previous findings suggested that HSP90α may be also regulating YAP activation through the modulation of Src/Yes1 kinases (**Fig. 3j**). Following a similar approach, we observed that *Hsp90aa1* silencing did not affect *Src* mRNA levels, but significantly downregulated *Yes1* in CAFs (**Fig. 4h**) and TGFβ1-stimulated NFs (**Fig. S8l**). Analyses of protein levels showed that depletion of *Hsp90aa1* in CAFs and TGFβ1-stimulated NFs had no effect on total Src (**Fig. 4i and S8m**). On the contrary, *Hps90aa1* RNAi treatment decreased the protein levels of Yes1. Furthermore, pulse-chase analysis with CHX showed that Yes1 protein stability was significantly reduced after *Hsp90aa1* depletion in CAFs (**Fig S8n**). Both Src and Yes1 can activate the transcriptional activity of YAP by promoting phosphorylation at Y357^33, 37^. Using a system were human YAP-YFP was constitutively expressed in NF^WT^/NF^KO^, which enabled the analysis of Y357 phosphorylation, we observed that *Hsp90aa1* depletion reduced YAP1 activation albeit it was not significant for a small margin (**Fig. 4j**). On the contrary, the strongest effect was still at the level of protein expression.

Altogether, these results indicate that HSP90α stabilizes YAP/TAZ protein levels and potentiates their activity. In conditions where YAP/TAZ are aberrantly activated, such as CAFs and TGFβ-stimulated NFs^12^, inhibiting HSP90α significantly diminishes their function. As TAZ has very limited functional relevance in CAFs^12^, our results suggest that YAP may be a key effector of HSP90α in controlling the pro-tumour activities in CAFs.

### HSP90α potentiates mechanotransduction in CAFs and constitutive expression of YAP attenuates deficiencies associated to *Hsp90aa1* depletion

YAP activity upregulates key cytoskeletal components such as myosin light chain 2 (MLC2), ANLN and DIAPH3 that promote the formation of stress fibres and focal adhesions (FAs) in CAFs, and subsequent mechanotransduction activation. These YAP-dependent changes promote the generation of aggressive CAF phenotypes with increased pro-tumour activities^10, 12, 13^. Immunoblot and immunofluorescence analyses demonstrated that *Hsp90aa1* silencing in CAFs reduced phosphorylation of MLC2, FAK and Paxillin (**Fig. 5a**). These pathways have been shown to affect the coupling between the actomyosin network and the ECM, increasing the number and size of FAs and the ability of cells to generate forces^38^. Morphometric analyses of FAs showed that *Hsp90aa1* silencing had marginal effects FA size but significantly reduced FA number and area per cell (**Fig. 5b&c**). To examine the particular role of the HSP90α:YAP axis in these phenotypes, we constitutively expressed YAP-YFP in NF^KO^, which led to a recovery of YAP expression levels (**Fig. 5d**). This was accompanied by a significant increase in FA size and number (**Fig. 5e&f**) and a recovery in the ability of NF^KO^ cells to remodel collagen gels (**Fig. 5g**). Overall, these results indicate a key function of HSP90α as a key upstream regulator of YAP in CAFs.

**Figure 5.**
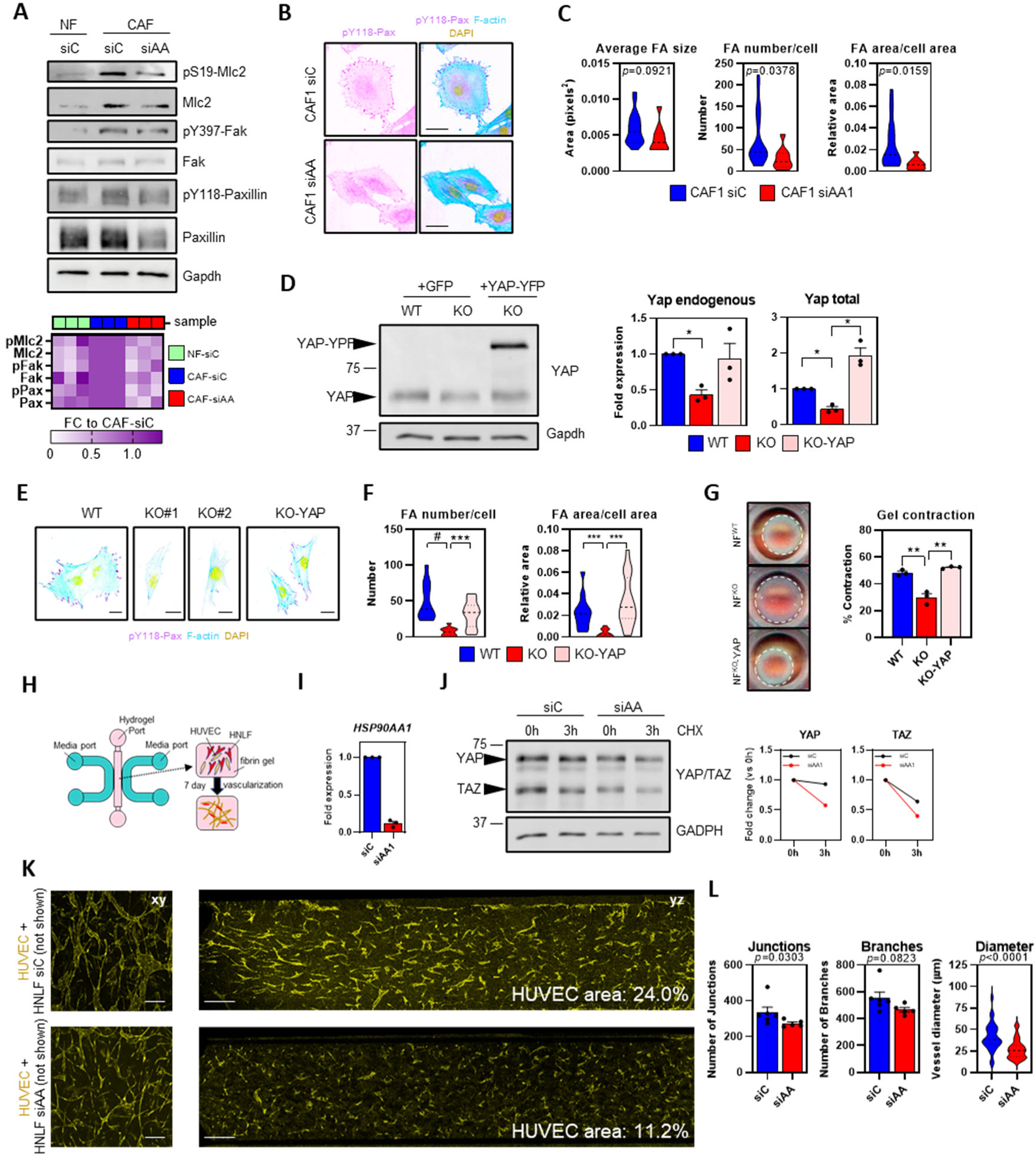
HSP90a potentiates mechanotransduction pathways in CAFs and constitutive expression of YAP attenuates deficiencies associated to Hsp90aal depletion. **A.** Western blots show expression of Gapdh, Mlc2, Fak and Paxillin, and corresponding phosphorylated (active) forms: phospho-Mlc2 (S19), phospho-Fak (Y397) and phospho-Paxillin (Y118) in NF1 and CAF1 after transfection with control (siC) and *Hsp90aal* (siAA) RNAi. Heatmap shows fold change (FC) expression values for each protein normalized to Gapdh against CAF-siC conditions (n=3). **B.** Images represent phospho-Paxillin (pY118-Pax; purple), F-actin (cyan) and DAPI (yellow) staining of CAF1 after transfection with control (siC) and *Hsp90aal* (siAA). Scale bar, 50 pm. **C.** Graphs show average size, number per cell and area per cell of FAs as inferred from images in (B). (CAF-siC, n=20; CAF-siAA, n=10 for all graphs). **D.** Western blots show expression of YAP and Gapdh in NF"^1^ and NF^KO^. Where indicated, NF^K0^ cells were stably expressing GFP or human wild­type YAP-YFP. Histograms show fold expression levels of endogenous and total YAP for each condition (n=3). **E.** Images show phospho-Paxillin (pY118-Pax; purple), F-actin (cyan) and DAPI (yellow) staining of NF^WT^, NF^K0^ (two different cells #1 and#2) and NF^K0^ expressing human wild-type YAP. Scale bar, 25 pm. **F.** Graphs show number per cell and area per cell of FAs as inferred from images in (E). (NF^WT^, n=13; NF^KO^, n=12; NF^KO^-YAP, n=17). **G.** Images show representative gel contraction assays for NF"^1^, NF^KO^ and NF^K0^ expressing wild-type YAP (NF^KO^-YAP). Graphs show percentage of gel contraction (n=3). **H.** Schematic illustrating the microfluidic set­up for analysis of fibroblast-dependent emergent vascularization. HUVECs and HNLFs are embedded in fibrin and submitted to flow for 7 days. **I.** Graph shows fold expression of *HSP90AA1* (relative to *GAPDH)* in HNLF after transfection with control (siC) or *HSP90AA1* (siAA) siRNAs (n=3). **J.** Western blot showing YAP/TAZ and GAPDH expression in HNLF after transfection with control (siC) or *HSP90AA1* (siAA) siRNAs (smart-pools) during pulse-chase with CHX at 0 h and 3 h. Graphs represents quantification of the indicated blots (relative to GAPDH) at 3 h of CHX treatment relative to the expression at 0 h for both conditions. **K.** Images show a representative confocal plane (left panels, xy) and a z-stack (right panels, yz) of anti-CD31 staining (HUVEC, yellow) of vascularization assays generated with HNLF after transfection with control (siC) or *HSP90AA1* (siAA) siRNAs. Fibroblasts are not shown. Area covered by HUVECs in each condition is shown. Scale bars, 200 pm. Graphs show number of junctions, branches and vessel diameter of HUVEC networks for each condition. (Junctions/Branches: HLNF-siC, n=6; HLNF-siAA, n=5. Diameter, n=30).

Next, we investigated if HSP90α-dependent YAP modulation was also relevant in other non-tumoral systems. Using microfluidic devices of endothelial cells (*i.e.* Human umbilical vein endothelial cells, HUVECs) and human normal lung fibroblasts (HNLF) to model emergent vascularization (**Fig. 5h**), we have recently shown that YAP-mediated mechanotransduction is required in fibroblasts to promote ECM remodelling and provide mechanical support for vascularization^39^. Transient knocked-down of *HSP90AA1* in HNLF (**Fig. 5i**) negatively affected YAP/TAZ stability in this system (**Fig. 5j**) and resulted in defective vascular networks with less interconnections and vessel diameter (**Fig. 5k&l**). Taken together, these results indicate that HSP90α participates in mechanotransduction programs by stabilizing YAP protein levels and sustaining their activity and function.

### Deletion of host *Hsp90aa1* inhibits tumour growth

YAP signalling and mechanotransduction pathways in CAFs engage in a positive feedback loop during BC progression, by promoting ECM remodelling, as well as cancer cell invasion and growth^10, 12, 13^. Having shown that HSP90α is a strong positive regulator of YAP signalling in CAFs, we investigated whether stromal HSP90α loss might alter the TME and subsequently affect BC pathogenesis. To test this, we performed orthotopic injections using the *Hsp90aa1* knock-out model. As previously reported^40^, ablation of *Hsp90aa1* was well tolerated and females had no observable phenotype (***not shown***). Isolated mammary fibroblast from *Hsp90aa1* knock-out adult females presented decreased levels of HSP90α mRNA and protein when compared to wild-type littermates, whereas HSP90β levels were not affected (**Fig. S3c&e**). Syngeneic E0771 murine BC cells (wild type for both *Hsp90aa1* and *Hsp90ab1*) were orthotopically injected into the mammary fat pad of immunocompetent wild-type (*Hsp90aa1^+/+^*), heterozygous (*Hsp90aa1^+/-^*) and knock-out (*Hsp90aa1^-/-^*) mice (**Fig. 6a**). A pronounced delay in tumour growth was observed in *Hsp90aa1^+/-^* and *Hsp90aa1^-/-^* mice compared to *Hsp90aa1^+/+^* littermates (**Fig. 6b**). Endpoint analyses of tumour volume and weight informed of significant decreases in *Hsp90aa1^-/-^* vs *Hsp90aa1^+/+^*mice (**Fig. 6c and Fig. S9a**).

**Figure 6.**
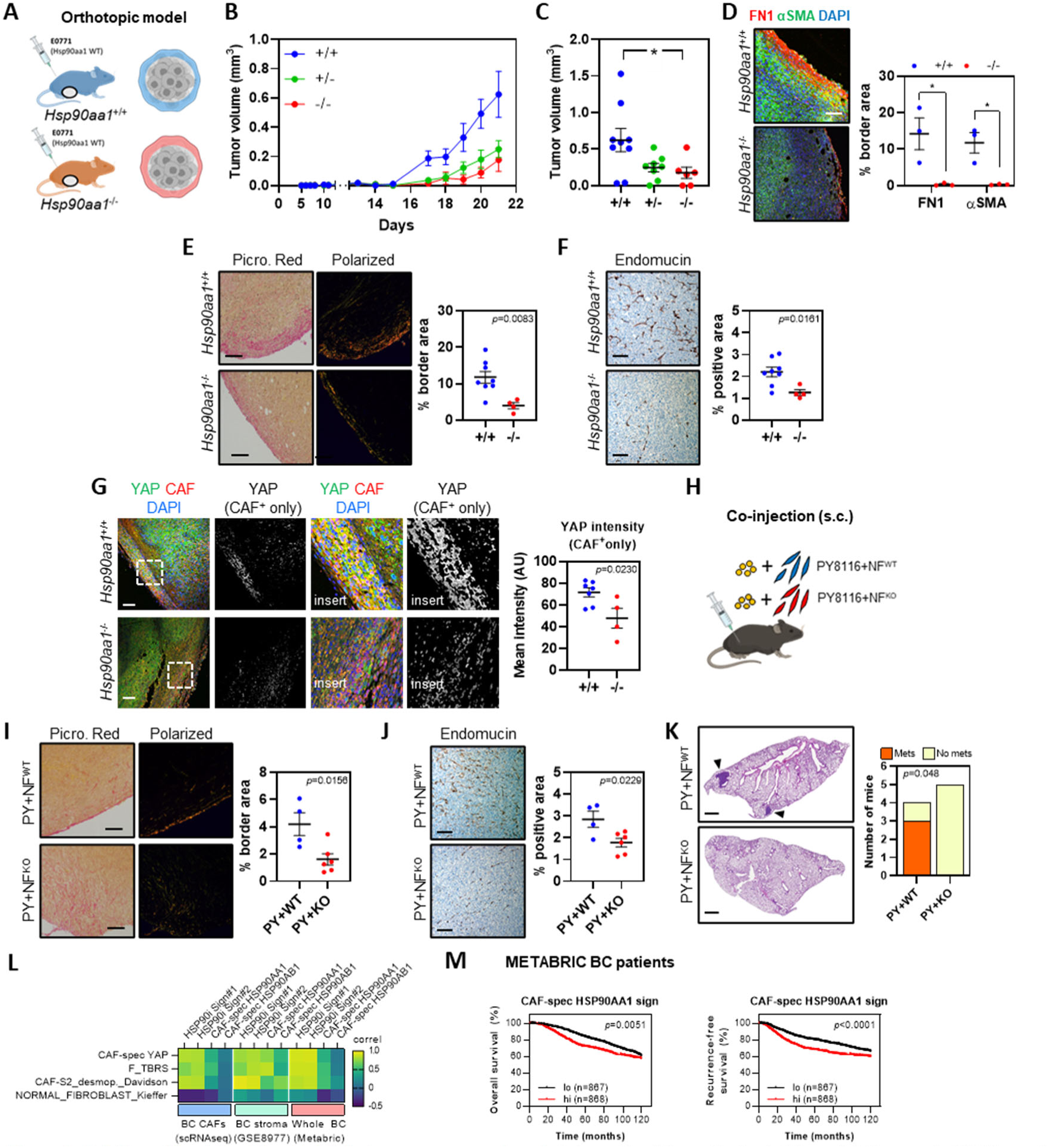
Deletion of host *Hsp90aal* inhibits tumour progression. **A.** Schematic of the orthotopic BC model. Syngeneic E0771 murine BC cells were injected on the mammary fat pad of *Hsp90*^/t^, HspdO^’* and *HspgO^* adult females (BL6 background, immunocompetent). **B.** Graph shows tumour growth curves of E0771 tumours growing in *HspdO*^, Hsp90^’* and *Hsp90’^/^’* mice *(Hsp90*^,* n=9; *Hspd^’,* n=8; *HspSO^,* n=6). **C.** Graph shows tumour volume at day 21 from tumours in (B). **D.** Images show Fnl (red), aSMA (green) and DAPI (blue) staining in sections from primary E0771 tumours (tumour border regions) grown in *Hsp90*^f^\** and *Hsp90^* mice. Scale bar, 100 pm. Graph on the right shows the percentage of border area positive for Fnl and crSMAin representative tumour sections (n=3). **E.** Images show picrosirus red staining (left) and polarized light from the same regions (right, representing fibrillary collagen) in sections from primary E0771 tumours (tumour border regions) grown in *Hspgo*^* and *Hsp90^* mice. Scale bar, 100 pm. Graph on the right shows the percentage of border area positive for fibrillary collagen (+/+, n=8; n=4). **F.** Images show endomucin staining in sections from primary orthotopic E0771 tumours grown in *Hsp90*^* and *Hsp90’^/^’* mice. Scale bar, 100 pm. Graph on the right shows the percentage of area positive for endumucin (+/+, n=8; n=4). **G.** Images show total YAP (green), S100A4 (CAF marker, red) and DAPI (blue) staining in sections from primary E0771 tumours (tumour border regions) grown in *Hsp9O*^* and *Hsp90’^/^’* mice. Scale bar, 100 pm. Greyscale panels represent processed images showing YAP staining in CAFs. Panels on the right show indicated zoom up areas (insert). Chart shows mean YAP intensity in CAFs (+/+, n=7; n=4). **H.** Diagram illustrating the approach for analysis of impact of fibroblast-specific depletion of *Hsp90aal* in tumour growth by subcutaneous implantation of metastatic MMTV-PyMT murine cancer cells (PY8116, BL6 background) together with NF^WT^ and NF^K0^ cells. **I.** Images show picrosirus red staining (left) and polarized light from the same regions (right, representing fibrillary collagen) in sections from primary PY8116+NF^WT^ (PY+NF^WT^) or PY8116+NF^KO^ (PY+NF^K0^) tumours grown in nude mice. Scale bar, 100 pm. Graph on the right shows the percentage of border area positive for fibrillary collagen (PY+NF^WT^, n=4; PY+NF^K0^, n=6). **J.** Images show endomucin staining in sections from primary PY8116+NF^WT^ (PY+NF^WT^) or PY8116+NF^KO^ (PY+NF^K0^) tumours grown in nude mice. Scale bar, 100 pm. Graph on the right shows the percentage of area positive for endumucin (PY+NF^WT^, n=4; PY+NF^K0^, n=6). **K.** Images show H&E staining of sections from lungs from nude mice growing PY+NF^WT^ or PY+NF^K0^ subcutaneous tumours, showing the presence of spontaneous lung metastases (arrowheads). Scale bar, 1 mm. Graph shows the number of mice growing PY+NF^WT^ or PY+NF^K0^ subcutaneous tumours that present evidence of lung metastases or not. *P* value for Fisher’s Exact Test is shown. **L.** Heatmap showing correlation coefficients between gene signatures for: YAP activation in CAFs ("CAF-spec YAP"), TGFp responses in fibroblasts ("F_TBRS"), desmoplastic CAF subtypes ("CAF-S2_desmop._Davidson") and normal fibroblast subtypes ("Normal_Fibroblast_Kieffer"), against HSP90i signatures (#1 and #2) and gene expression signatures for HSP90AA1 and HSP90AB1 in CAFs. Correlation analysis was performed in BC CAF datasets (extracted from scRNAseq^23^), BC stroma datasets (GSE8977) and whole BC tumours (Metabric dataset). **M.** Graphs show percentage of overall survival (left) and recurrence-free survival (right) of BC patients from the Metabric dataset with high (hi) or low (Io) expression of a gene signature associated with HSP90AA1 activity in CAFs. Where indicated, individual *p* values are shown; alternatively, the following symbols were used to describe statistical significance

A dense fibrillary collagen network is associated with increased tissue stiffness, which promotes tumour progression and dissemination in BC^9, 41^. We evaluated if ablation of host *Hsp90aa1* was linked to defects in ECM and thus in desmoplastic CAF activity. Immunofluorescence approaches showed decreased fibronectin (Fn1, an ECM component) and smooth muscle actin (αSMA, a marker of CAF/myCAF) in tumours from *Hsp90aa^-/-^* mice when compared to tumours from wild-type littermates (**Fig. 6d**). In addition, immunohistological analyses of tumours showed reduced levels of collagen deposition/desmoplasia in *Hsp90aa1^-/-^* compared to *Hsp90aa1^+/+^* mice, as read by picrosirius red (**Fig. 6e**) and Masson’s trichrome stainings (**Fig. S9b**). YAP activity in CAFs can also promote angiogenesis in vivo^10, 12, 13^. In agreement, we observed a significant reduction in blood vessel coverage in tumours grown in *Hsp90aa1^-/-^* mice (**Fig. 6f**). Similar to our in vitro data, CAFs in knock-out mice showed reduced YAP staining, reinforcing the link between HSP90α, YAP and CAF functionality (**Fig. 6g**). As fibroblasts also play a crucial role in the establishment of the metastatic niche^42^, we speculated that *Hsp90aa1* deficiency could have an inhibitory effect on experimental lung metastasis (**Fig. S9c**). Tail vein injection of E0771 cells resulted in efficient lung colonization in *Hsp90aa1^+/+^* mice at 4 weeks, which was significantly reduced in *Hsp90aa1^-/-^* mice (**Fig. S9d**).

In these models *Hsp90aa1* is not solely silenced in fibroblasts, but also in other cell populations of the TME such as endothelial and immune cells. Conditional targeting of fibroblasts in mice is problematic, as available *cre* mouse models employed to target fibroblasts may also alter other cellular compartments^43^. To simplify the system and to ensure that only fibroblasts were depleted of *Hsp90aa1*, we proceeded to validate our findings by engrafting a MMTV-PyMT cancer cell line (PY8116 model) with NFs isolated from mammary glands of *Hsp90aa1^-/-^* or *Hsp90aa1^+/+^*mice (NF^WT^/NF^KO^ system)(**Fig. 6h**). Contrary to experiments with stromal *Hsp90aa1* deletion, the tumour growth curves did not show significant differences between the two groups in immunocompromised or immunocompetent mouse strains (**Fig. S9e&f**). However, immunohistochemistry analyses showed relevant differences in stromal tissue remodelling with decreased collagen deposition and angiogenesis in PY8116+NF^KO^ tumours when compared to PY8116+NF^WT^ (**Fig. 6i&j and Fig. S9g**). Contrary to E0771 model, the PY8116 cells spontaneously metastasize to the lung^44^, which enabled us to assess the role of fibroblast HSP90α in BC dissemination. Indeed, we observed overt spontaneous metastasis in the lungs of mice growing PY8116+NF^WT^ tumours that were completely absent in the lungs of mice growing PY8116+NF^KO^ tumours (**Fig. 6k**). Altogether, these findings demonstrate that HSP90α is a regulator of aggressive CAF phenotypes *in vivo* through YAP, and that perturbations in HSP90α can be detrimental on the generation of aggressive TMEs and metastatic spread.

### Association between HSP90AA1 activity in CAFs and disease progression in patients

To investigate the relevance of our findings in human malignancies, we analysed the expression of genes and gene signatures associated to YAP activity and CAF subtypes in relation to HSP90α activity. Since we did not observe significant changes in *HSP90AA1/HSP90AB1* gene expression but changes in associated gene expression programs between NF and CAFs, we generated a gene signature of the top 500 downregulated genes after HSP90α depletion in CAFs that were not affected by HSP90β as a surrogate for specific HSP90α activity in CAFs (‘CAF-specific HSP90AA1 signature’)(**Supplementary Table 2**). A similar approach was employed to generate ‘CAF-specific HSP90AB1 signature’. For YAP/TAZ activity we relied on a previously reported signature^10^. Using a human BC scRNAseq dataset^23^, we observed that CAF-specific HSP90AA1 and YAP signatures, as well as associated individual genes were preferentially expressed in CAFs from mCAF phenotypes in comparison to other subpopulations (**Fig. S9h&i**). In agreement, a strong positive correlation between HSP90i and CAF-specific HSP90AA1 signatures, and gene sets related to YAP activation, TGFβ responses and desmoplastic CAF subsets was detected (**Fig. 6l**). Conversely, the association was primarily negative with NF signatures. By contrast, the CAF-specific HSP90AB1 signature displayed no relevant correlations. To further probe this relationship, we investigated these associations using independent BC stroma and whole tumour gene expression datasets, observing similar results (**Fig. 6l and Fig. S9j**). Notably, high expression of CAF-specific HSP90AA1 signature was also significantly associated with poor overall and recurrence-free survival in BC patients from the METABRIC cohort (**Fig. 6m**). On the other hand, CAF-specific HSP90AB1 signatures presented marginal correlation with these parameters (**Fig. S9k**). Of note, the expression of the *bona fide* myCAF marker *ACTA2* (encoding for αSMA) presented no prognostic associations in this cohort (**Fig. S9m**), underlying the specific role of HSP90α. The prognostic value of HSP90α signatures was further validated in colon and ovarian cancer patient datasets (**Fig. S9n**).

Together, these findings suggest that HSP90α activity in CAFs dictates signalling and transcriptional programs implicated in CAF emergence and functions, promoting ECM remodelling to support the growth of primary tumours and metastatic dissemination that ultimately impact patient prognosis.

## DISCUSSION

Given the crucial role of CAFs in the TME and tumour progression, understanding the mechanisms governing their pro-tumour behaviour is essential for identifying vulnerabilities and developing new therapies. The response of TME components to stress is an emerging area of interest^7^, as stromal cells may develop targetable dependencies similar to cancer cells.

This study identifies HSP90α as a key regulator of CAF function, influencing ECM remodelling and organization as well as angiogenesis, tumour growth and dissemination in experimental models of BC. Mechanistically, *Hsp90aa1* deletion impaired cellular responses to TGFβ stimulation in NFs, preventing their transition to a myofibroblast phenotype. This finding aligns with previous studies showing that HSP90 stabilizes TGFβ receptors^25^, of particular relevance fibrotic contexts^17, 18^. In a tumoral setting, HSP90α may support CAF emergence during early tumour development, when inflammation and desmoplasia increase TGFβ levels. In agreement, our in vivo studies showed a reduction in αSMA-positive CAFs, fibronectin and collagen in tumours generated in *Hsp90aa1* knock-out mice. Additionally, HSP90α interacts with YAP, stabilizing its protein levels and ensuring the sustained mechanotransduction activity characteristic of CAFs. Other TFs aberrantly activated in CAFs such as β-catenin^10^ were not altered by *Hsp90aa1* depletion, underlying the specificity of HSP90α regulation over YAP. Restoring YAP levels rescued functional defects associated with HSP90α loss, highlighting its pivotal role in CAF functionality. While prior studies linked HSP90 to heat-induced YAP activation via the Hippo pathway^32^, our findings reveal a distinct mechanism, emphasizing a direct interaction between HSP90α and YAP stabilization. Importantly, this regulation is independent of heat-shock stress and appears particularly relevant in mechanically stressed contexts such as CAFs^12^, where mechanotransduction dominates over Hippo-dependent control. Further validation of this mechanism was achieved using fibroblast-dependent vascularization models. This underscores the broader relevance of the HSP90α-YAP axis beyond CAFs. Together, these findings represent an unprecedented layer of interaction between proteostasis and mechanotransduction control, and its impact in cancer and physiology.

Notably, HSP90β depletion had no significant effects, emphasizing the isoform-specific functions of HSP90 that need to be taken into account in future studies^14^. HSP90α is the main inducible isoform upregulated under stressful conditions^12^ and presents an extracellular function^18^. Moreover, both HSP90α and HSP90β form specific complexes with co-chaperones, modulating distinct sets of client proteins and participating in different signalling cascades. Contrary to tumour cells^45^, CAFs do not appear to upregulate *HSP90AA1* expression in vitro or in vivo, albeit a mild increase in protein levels was observed in CAF1 vs NF1. Given that the in vitro conditions did not impose extrinsic stress, we hypothesise that HSP90α may be essential in managing intrinsic stresses associated with the pathologically activated phenotype of CAFs, akin to mechanisms observed in mechanically stressed systems^46^. HSP90-dependent transcriptional programs are a consistent feature of CAFs, particularly enriched in myCAF phenotypes. myCAFs present a distinct phenotype characterized by excessive ECM production and a robust cytoskeleton^47^, which, if uncontrolled by proteostasis, may compromise cell viability. We propose that CAFs are particularly vulnerable to HSP90α loss because this chaperone stabilizes protein levels of key signalling nodes or TFs aberrantly activated in CAFs such as TGFβ-SMAD and YAP pathways.

In murine BC models, HSP90α silencing in the stroma or CAFs led to decreased fibrillary collagen, CAF activation, and angiogenesis – processes dependent on YAP activity^10, 12, 13^. Tumour growth and metastasis were also affected, with varying impacts depending on the specificity of HSP90α loss, suggesting additional defects in immune or endothelial cells that require further examination. BC patient data revealed that HSP90α-dependent transcriptional signatures correlated with YAP activity, desmoplastic responses in CAFs, and survival. These findings suggest that targeting HSP90α could alter CAF function with therapeutic implications that may also be extended to other pathologies characterized by aberrant YAP activity. While available HSP90 inhibitors affect YAP stability, they target both HSP90 isoforms, potentially leading to toxicity due to the critical role of HSP90β in homeostasis. Developing selective HSP90α inhibitors could provide a safer therapeutic strategy.

## METHODS

### Mouse strains

C57BL/6 *Hsp90aa1^+/+^* (wild-type), *Hsp90aa1^-/-^* (knock-out) and *Hsp90aa1^+/-^* (heterozygous) mice^40^, as well as CD-1 Nude mice (Charles River), were kept in a pathogen-free animal facility unit and used as recipients for tumours. All animals were kept in accordance with EU regulations under regional Animal Experimentation licenses PI/05/19 and PI/02/23.

### Cell lines

Established murine fibroblasts from FVB/n normal mammary glands (NF1, NF4) and MMTV-PyMT mammary carcinoma (CAF1, CAF3, CAF5) have been previously described^12, 48^. Murine mammary fibroblasts were also isolated and established from the healthy mammary glands of *Hsp90aa1^+/+^* and *Hsp90aa1^-/-^* mice (NF^WT^ and NF^KO^). Briefly, normal mammary glands were enzymatically digested (1% collagenase) and fibroblasts isolated and HPV-E6 immortalized as described^48^. Human primary fibroblasts from normal human lungs (HNLFs) were provided by Emad Moeendarbary (University College London, UK)^39^. This cell line was not immortalized and used at low passage (less than 10). All fibroblasts were cultured in DMEM high glucose (Sigma), GlutaMax (Gibco), 10% FBS, and Pen/Strep antibiotics (complete media) unless otherwise specified.

D2A1 (Cellosaurus CVCL_0I90) murine BC cells were derived from spontaneous mammary tumours in originated from a D2 hyperplastic alveolar nodule. These cells were a kind gift from Erik Sahai (Crick Institute, UK), and were used in *in vitro* experiments. E0771 (Cellosaurus CVCL_GR23) murine BC cells were originally isolated from a spontaneous tumour in C57BL/6 mouse. They were a kind gift from Kristian Pietras (Lund University, Sweden), and were used to generate syngeneic tumours in C57BL/6 mice. D2A1 and E0771 were cultured in complete media. PY8119 cells are murine BC cells isolated from the MMTV-PyMT (C57BL/6) model. They were a kind gift from the lab of Hector Peinado (CNIO, Spain) and originally isolated by Daniela Quail lab (Goodman Cancer Institute, Canada). These cells stably express luciferase and GFP (luc-GFP) and were cultured in Ham’s F12K + 5% FBS (unless stated otherwise) and employed in *in vitro* assays and to generate tumours in C57BL/6 mice.

Human Umbilical Vein Endothelial Cells (HUVECs) were cultured in EBM-2 Basal Medium (Ref. #CC-3156, Lonza) supplemented with EGMTM-2 MV SingleQuots™ Supplement Pack (Ref. #CC-4147, Lonza) in flasks/dishes pre-coated with Rat Tail Collagen I (#354249 Corning®) at a concentration of 0.05mg/mL.

All cell lines were grown in a standard humidified 5% CO2 incubator at 37° C and tested negative for mycoplasma infection with MycoAlert^TM^ (Lonza). Cancer cell lines were fluorescently labelled using pCSII-IRES2-blasti-eGFP.

### Generation of CRISPR knock-in fibroblast cell lines

NF1 and CAF1 lines were genetically modified to incorporate a V5-dTAG sequence in frame at the C-terminus of the *Yap1* locus (in heterozygosis). As a result, these cells produce a copy of the wild-type Yap1 and one copy of Yap1-V5-dTAG. The product of this modified version can therefore by detected and immunoprecipitated with V5 antibodies. The dTAG sequence is FKBP12^F36V^. Addition of the targeted degrader dTAG^V^-1 (dTAG) in these cell lines results in the targeted degradation of the chimeric Yap1-V5-dTAG, remaining the wild-type protein unaltered. The sgRNA sequences including PAM region targeting exon 9 of *Yap1* was determined using UCSC Genome Browser as follows: 5’-CACGTGGTTATAGAGCTGCAGGG-3’. sgRNA oligonucleotides were cloned into pX330A-sgX-sgPITCh at BsbI restriction site. The design of the repair vector was performed as it was previously described^49^.

The left microhomology sequence 5’-TCTCACGTGGTTAGGAAGCT-3’ and right microhomology sequence 5’-CAGACTCAGAGTGGCTCCCTGC-3’ were cloned into vector pCRIS-PITChv2-C-dTAG-BSD. This vector was previously modified for the expression of the V5 epitope and neomycin resistance. Fibroblasts were transfected with both plasmids using Lipofectamine (Life Technologies) following manufacturer’s instructions. After 48 h, G418 (750 µg/mL) was added to the media for the selection of the positive cells, and 5 days later cells were single sorted into a 96 well plate. Individual clones were expanded and the YAP1 locus targeted by CRISPR was sequenced for knock-in validation. In addition, the Yap1-V5-dTAG and endogenous proteins expression were also confirmed by immunoblotting.

### cDNA, RNAi and reagents

siRNAs were purchased from Dharmacon and are listed in the **Supplementary Table 3**. Lentiviral plasmids for expression of wild-type and mutant YAP were YAP1-YFP (#112284, Addgene) and S5A-YAP1-YFP vectors (#112285, Addgene), respectively. These plasmids were a kind gift of Erik Sahai (Crick Institute, UK). For *in vitro* treatments, the following growth factors and drugs were used: recombinant human TGFβ1 (#100-21C-10UG, Peprotech), Geldanamycin (#S2713, Selleckchem), Tanespimycin/17-AAG (#S1141, Selleckchem), Cycloheximide/CHX (C7698, Sigma), dTAG^V^-1 (#6914, Tocris), DMSO (#D8418, Sigma), L-Ascorbic Acid (#A92902, Sigma), Mitomycin-C from *Streptomyces caespitosus* (#M4287-2G, Sigma), SB431542 (#616461, Sigma).

### Animal experiments

#### Generation of syngeneic orthotopic tumours

E0771 murine BC cells were used to generate orthotopic tumours in the mammary gland of C57BL/6 *Hsp90aa1^+/+^*, *Hsp90aa1^-/-^* and *Hsp90aa1^+/-^* female mice. Briefly, 2.5 × 10^5^ E0771 cells were injected directly into the 4^th^ mammary fat pad of 6-8 weeks old mice by performing small surgery to expose the mammary gland, that was then closed with suture. Tumour growth was monitored every other day using palpation and callipers. Tumour volume was calculated for each time point using the formula: (4π/3)*x*(width/2)^2*x*(length/2). When the end point was reached, mice were sacrificed and tumours were extracted, weighted and processed.

#### Generation of experimental metastases

E0771 murine BC cells were used to generate lung metastasis of C57BL/6 *Hsp90aa1^+/+^*, *Hsp90aa1^-/-^* and *Hsp90aa1^+/-^* female mice. Briefly, 2.5 × 10^5^ E0771 BC cells were injected via the lateral tail vein. Mice weight was monitored every two days until weight loss started. Mice were sacrificed approximately 4 weeks after the procedure, when no more than 20% of their weight was lost. Lungs were then extracted and processed.

#### Generation of subcutaneous tumours

PY8119 murine BC cells co-injected with NF^WT^ or NF^KO^ were used to generate subcutaneous tumours in female mice. Briefly, 5 × 10^5^ PY8119-luc-GFP cells and 1.5 × 10^6^ NFs were suspended in 100 μL of PBS:Matrigel (50:50) and injected subcutaneously into 6-8 week old wild-type CD-1 nude or C57/BL6 mice. Tumours were monitored and processed as with orthotopic tumours. In addition, lungs were also acquired and further processed for spontaneous metastasis analyses.

### Transfections and cell perturbations

Fibroblasts were seeded at 60% confluence and transfected using DharmaFECT 1 (Dharmacon) with siRNA (100 nM final concentration) following manufacturer’s instructions. Fibroblasts were used 24 h or 48 h after transfection for subsequent cellular or molecular analysis. siRNA description can be found in **Supplementary Table 3**. TGFβ1 was added to a final concentration of 5 ng/mL into the cell culture media 5 h after transfection, and the stimuli was maintained throughout the whole experiment. Cells were typically used after 24 h of TGFβ1 stimulation. Cell lines stably expressing cDNA (YAP^WT/MUT^-YFP or GFP) were generated by lentiviral infection. Fluorescent-labelled cells were sorted by FACS. TGFβ receptor inhibitor SB431542 was used at a final concentration of 5 μM for 24 h. Geldanamycin and Tanespimycin were used at 10 nM concentration for 24 h. dTAG^V^-1 was used at 300nM for 24 h to degrade Yap1-V5-dTAG. For protein stability assays, cells were treated with 10 µg/mL of CHX for the indicated intervals

### ECM-remodelling assay

To assess force-mediated matrix remodelling, 10^5^ murine NF/CAFs were embedded in 100 µL of a mixture of collagen I (BD Biosciences), ECM gel mixture (Sigma), 10% FBS and DMEM yielding a final collagen concentration of approximately 4.6 mg/mL and a final ECM gel mix concentration of approximately 2.2 mg/mL in 48-well plates precoated with 5% BSA. Once the gel was set, cells were maintained in complete medium with any relevant factor/inhibitor added. Gel contraction was monitored daily by scanning the plates. Unless stated otherwise, the gel contraction value refers to the contraction observed after 2 days. To obtain the gel contraction value, the relative diameter of the well and the gel were measured using ImageJ software, and the percentage of contraction was calculated using the formula 100 × (well area – gel area) / well area.

### Cell Derived Matrices (CDM) experiments

#### Generation of CDMs

CDMs were generated as described^50^. Briefly, glass-bottom 12-well plates were coated with 0.2% (w/v) gelatine o/n at 37°C and cross-linked with 1% (v/v) Glutaraldehyde for 1 h. After washing with PBS, 1 M Glycine was added for 1 h, and washed with PBS. Then, fibroblasts were seeded into a confluent monolayer and culture media was supplemented with 5 ng/mL of TGFβ1 and 50 µg/mL ascorbic acid to promote ECM deposition. Cells were then maintained in culture for 5 days and media was changed every other day. For the experiments without fibroblast removal, plates were directly fixed with 4% paraformaldehyde (PFA) for 30 min at RT and further processed for immunofluorescence analyses. For experiments with fibroblast removal, cells were lysed and the resulting CDMs were used right away or stored with 1 mL of PBS at 4 °C until further use.

#### Cancer cells cultured over CDMs

5 × 10^3^ D2A1-GFP cells were seeded on top of CDMs in low serum media (2% FBS) and maintained in culture for 4 days. To monitor cancer cell growth, GFP-fluorescence images were taken every day with an Eclipse TS100 (Nikon) microscope. Plates were fixed in 4% PFA and processed for IF.

#### Cancer cell spheroids time-lapse over CDMs

D2A1-GFP spheroids were generated by plating 100 cells per well in a low-adherence curved-bottom 96 well plate for 48 h. Then, 4 spheroids per condition were transferred over CDMs in complete cell culture medium. Images were taken every hour for 3 days with a Nikon Eclipse Ci2 time-lapse microscope coupled to an environmental chamber.

#### CDM-based analyses

To measure CDM structural parameters, the TWOMBLI Fiji macro was employed^51^. To analyse cancer cell behaviour parameters, GFP positive areas were identified and the ‘Analyze particles’ tool from Fiji was used to measure the distribution of angles in which the cancer cells were disposed on top of the CDMs generated by fibroblasts. In addition, total GFP area was calculated for each image and CDM parameters were calculated as described before. For the analysis of clusters, the area and shape features of endpoint clusters were analysed using Fiji.

### Transwell assay

1 × 10^4^ fibroblasts were seeded on top of 8 μm pore inserts (24 well plate/12 insert chamber, #3422, Costar) in complete medium. The lower chamber was filled with complete media. In the case of the TGFβ1-stimulated conditions, the total concentration in the well (upper and lower chamber together) was 5 ng/mL. After 24 h, media was aspirated, and cells were removed from the upper part of the membrane. Next, membranes were stained using Crystal Violet staining solution for 20 min, washed with H_2_O and dried o/n. The following day, membranes were imaged with an Eclipse TS100 (Nikon) microscope with the 4× objective and analysed with Image J.

### Cancer cell:fibroblast co-culture experiments

#### 2D co-cultures

3 × 10^5^ fibroblasts were cultured to a confluent monolayer in 6-well plates. The next day, 10 μg/mL of Mitomycin-C was added for 3 h. Cells were washed with PBS and cultured with either DMEM supplemented with low serum media (2.5% FBS) or low glucose DMEM media supplemented with 2% FBS. On top of the fibroblasts, 1.5 × 10^4^ D2A1-GFP or PY8119-luc-GFP cells were seeded per well. Cancer cell growth was monitored by taking images in the fluorescent microscope every 24 h up to 7 days. Images were then analysed using Fiji to calculate cancer cell area.

### 3D co-cultures

24-well MatTek glass bottom plates were pre-coated with 150 µL of a mix of 1:1 Growth Factor Reduced Matrigel:PBS. Next, 1.5 × 10^4^ fibroblasts and 5 × 10^3^ D2A1-GFP cells were seeded on top in low glucose DMEM supplemented with 2.5% FBS. Cancer cell growth was monitored by taking images in the fluorescent microscope every 24 h up to 3 days. Images were then analysed using Fiji to calculate cancer cell area.

### Cell cycle analyses

Transfected fibroblasts were fixed in 70% ethanol o/n at 4°C 24 h after transfection. After fixation, cells were collected, washed once with PBS 1X, and suspended in 100 μL of PBS 1X containing 100 μg/mL RNAse A and 10 μg/mL Propidium Iodide. After 30 min of incubation, cells were classified at different cell cycle stage based on DNA content using flow cytometry.

### Emergent vascular morphogenesis assay

This was performed with 3-channel devices as described previously^39^. The device consists of three adjacent, parallel channels having rectangular cross sections with specific dimensions^39^. Adjacent channels are separated by continuous polydimethylsiloxane (PDMS) lips to contain the injected cell-embedded pre-polymerized fibrin gel. Acrylic moulds were fabricated by cutting (Epilog laser cutter) the designs through 1-mm-thick acrylic sheets. In three-channel devices, 0.3-mm-deep grooves were etched on the top surface of the acrylic sheet. Next, the laser-cut acrylic geometries were bonded to larger acrylic bases. Melds were then taped to the bottom of 60-mm Petri dishes to fabricate the final PDMS devices. Briefly, PDMS and cross-linker (10:1; SYLGARD 184; Dow Corning) were mixed, degassed, and poured over the mild to a height of 6 mm. After curing for 4 h at 60°C, the PDMS was cut and peeled from the mild. Gel filling ports and medium access ports were created by punching out 1- and 4-mm holes, respectively, using a biopsy punch (Miltex). The device was cleaned with tape to remove debris and then sterilized in an autoclave. Last, the bottom of the device and a glass coverslip (No. 1; Electron Microscopy Sciences) were treated with plasma for 60 s and pressed firmly together to form an irreversible bond. To restore hydrophobicity of surfaces and create irreversible bonding, the final devices were placed inside an 80°C oven o/n. HUVECs and human lung fibroblasts were suspended in EGM-2MV and mixed in 100 μL of a final encapsulation suspension containing HUVECs (4 million cells/mL) and fibroblasts (2 × 10^6^ cells/mL) in fibrinogen (3 mg/mL, Sigma-Aldrich) with thrombin (2 U/mL, Sigma-Aldrich). The suspension was mixed slowly several times over ice to avoid premature polymerization and then pipetted into the vascularization channel, *i.e.* half the volume through each of the two gel filling ports. The device was immediately placed in a humidified chamber and incubated at RT for 30 min for fibrin polymerization. Last, the medium channels were filled with warm EGM-2MV through the medium access ports. The devices were incubated at 37°C and 5% CO2, and the growth medium in the medium channels was replaced every day. On day 6, media was removed and EGM-2MV containing labelled anti-CD31 (REF #561654 Alexa Fluor® 647 Mouse Anti-Human CD31, BD Pharmingen) was added o/n to label HUVECs. The following day, media was replaced and images acquired in a Leica TS6 confocal microscope. The acquired images were analysed using ImageJ as described^39^.

### Immunohistochemistry (IHC) and immunofluorescence (IF)

#### Tissues

For histological analysis, tumours were dissected and fixed in 4% PFA for 24 h, dehydrated through a series of graded alcohols, and embedded in paraffin. Sections of 4 µm thickness were de-paraffined and rehydrated. Samples were processed for Haematoxylin-Eosin, Masson’s Trichrome or Picrosirius Red staining following standard procedures. For IHC, endogenous peroxidase was blocked by incubating sections in a 2% H_2_O_2_ dilution in methanol. After antigen retrieval using pH 6 sodium citrate buffer and blocking all non-specific binding sites, sections were incubated o/n at 4°C with primary antibodies. Sections were incubated with the secondary antibody for 30 min, followed by Streptavidin-HRP (ab64269, Abcam) for 1 h at RT. After performing the avidin-biotin complex technique, they were treated with metal-enhanced 3,30-diaminobenzidine (34065, Thermo Scientific) and counterstained with diluted haematoxylin. Antibody description and working dilutions can be found in **Supplementary Table 4**. Glass slides were scanned using Axio Scan.Z1 Slide Scanner (ZEISS).

For samples stained with picrosirius red, sections were serially imaged using with an analyser and polarizer oriented parallel and orthogonal to each other. Microscope conditions (lamp brightness, condenser opening, objective, zoom, exposure time, and gain parameters) were constant throughout image acquisition.

For IF analysis of tissues, after antigen retrieval sections were permeabilized with 0.2% Triton X-100 and incubated with 1% Goat serum (016201, Invitrogen) 5% BSA blocking solution for 30 min at RT. Sections were incubated overnight at 4°C with primary antibodies diluted in PBS. After washing, sections were incubated with fluorescent-conjugated secondary antibodies (AlexaFluor) and DAPI diluted in PBS for 1 h at RT. Before mounting, samples were treated for 10 min with 10 mM CuSO₄ / 50 mM NH₄Cl solution to eliminate paraffin autofluorescence. Confocal images were captured using a Leica SP5 microscope (20× objective). Antibody description and working dilutions can be found in **Supplementary Table 4**.

IHC and IF images were analysed using QuPath and ImageJ, and data analysed in GraphPad Prism. For each staining, positive area was calculated and normalized to the total area (or indicated regions of interest) for all images processed. Metastatic index was calculated as the area covered by metastases normalized to total area of the lung). Quantitative analysis of YAP staining in the tumour stroma was performed using ImageJ. Briefly, fibroblasts were identified using automated threshold based on S100A4 staining on stromal rich areas (tumour border). After background correction, the mean intensities of YAP staining were then measured. In all cases, all images from individual tumours were averaged for a final score per tumour.

#### Cells and CDMs

Cells were fixed in 4% PFA for 10 min and permeabilized by incubation in PBS 0.2% Triton 100 (Sigma) at RT for 7 min. Samples were blocked for 60 min at RT in blocking solution: 4% BSA PBS 0.05% Tween20 (Sigma). Then, cells were incubated with primary antibody in blocking solution in a wet chamber overnight at 4°C. After 3 washes of 15 min in PBS, secondary antibody in blocking solution was added for 1 h. After 3 washes of 15 min in PBS, samples were mounted and imaged using a Leica SP6 confocal microscope. CDMs were permeabilizated with a 0.5% PBS-Triton solution for 10 min at RT and then blocked o/n at RT with 3% BSA. Next, CDMs were incubated with PBS 0.1% Tween for few seconds prior incubation with the primary antibody for 3 h at RT followed by an incubation with the secondary antibody for 1h at RT. CDMs were imaged as described for cells. Unless stated otherwise, all acquisition settings were kept constant for all images, and all independent replicates imaged on the same session. Antibody description and working dilutions can be found in **Supplementary Table 4**.

Analyses of CDMs has been previously explained. For analyses of FAs, images of cells were analysed using ImageJ. The total area of individual cells was identified and measured using the phalloidin staining. Next, the same area was analysed to determine the FA parameters based on pY118-Paxillin staining. A specific threshold was applied in 8-bit converted images and the average size and total area of pY118-Paxillin positive regions was obtained. All images were subjected to the same threshold parameters to obtain the positive regions, and regions with areas smaller than 10 square pixels were discarded. The pY118-Paxillin-positive area relative to the whole cell area, number of pY118-Paxillin-positive areas and their average size was then calculated for each individual cell.

### Co-Immunoprecipitation

Cells were lysed in in immunoprecipitation lysis buffer (1% Triton X-100, 20 mM Tris-HCl pH 7.5, 50 mM NaCl, 50 mM NaF, 15 mM Na_4_P_2_O_7,_ 2 mM Na_3_VO_4_) containing protease and phosphatase inhibitors. Lysates were pre-cleared using G-sepharose, incubated with anti-V5 or anti-HSP90 antibodies (#1) or IgG immobilized on G-sepharose beads at 4°C for 4 h. Beads were then washed 3 times in washing buffer (50 mM Tris-HCl pH 7.5, 150 mM NaCl, 5% Glycerol, 0.05% NP-40) and eluted with 2X Laemmli Buffer.

### Luciferase reporter assays

The following plasmids were used for luciferase reporter assays: TEAD-reporter construct (pGL3-5xMCAT(SV)-49, that consists of 5 repeats of a TEAD binding element upstream of a minimal SV40 promoter that drives expression of the gene that encodes Firefly Luciferase.) and pEGFP-empty. CAFs were seeded in 6-wells plates at 70% confluency. Cells were transfected with the TEAD-reporter and pEGFP constructs using Lipofectamine 3000 according to the manufacturer’s instructions. Cells were lysed 48 h after transfection in 100 μL of lysis buffer (Promega). Aliquots of the cell lysates were used to read luciferase emission using Dual-Glo® Luciferase Assay System (Promega) according to the manufacturer’s instructions. A Western blot against GFP expression was performed with the same lysates. Firefly luciferase activities were normalised to GFP expression. Fold changes against control conditions were calculated.

### Western Blotting (WB)

Unless stated otherwise, cells were lysed directly in 2× Laemmli Buffer, sonicated and boiled. Alternatively, cells were lysed in RIPA buffer, mixed with Laemmli Buffer and boiled. Protein lysates and immunoprecipitants were processed following standard procedures for SDS-PAGE electrophoresis. Nitrocellulose membranes were blocked in BSA and incubated with primary and secondary antibodies. Dilutions and references for antibodies can be found in **Supplementary Table 4**. Unless described otherwise, HSP90 immunoblotting was performed with antibody #1. For protein detection, the Enhanced Chemoluminiscence (ECL) reaction was performed, and proteins were detected using the Amersham ImageQuant 800 (Cytiva). Exposures within the dynamic range were quantified by densitometry using ImageJ. For protein stability assays, cells were treated with 10 µg/mL of cycloheximide (CHX) for the indicated intervals and then lysed and subjected to WB. Immunoblot images in Figures show molecular weight markers in kDa.

### RNA isolation and qRT-PCR

RNA was extracted with the NZY Total Isolation Kit (NZYTech), according to manufacturer’s instructions. Reverse transcription was performed using the NZY First-Strand cDNA Synthesis Kit (NZYTech) and RT-qPCR using NZYSpeedy qPCR Green Master Mix 2X ROX plus (NZYTech) in a StepOnePlus real-time PCR system (Applied Biosystems). *Gapdh*/GAPDH was used as the housekeeping gene for the normalization of the data, and the quantification and comparison were obtained using the ΔΔCt method. Primers used and their corresponding sequence are described in **Supplementary Table 5**.

### RNA-sequencing processing and analysis

RNA was extracted as previously explained and checked by RNA ScreenTape analysis in an Agilent TapeStation system (Agilent Technologies). Samples were then sent to the *Centro Nacional de Análisis Genómico* (CNAG) for RNA sequencing analysis. Samples were first quality checked and transformed into a library of stranded mRNA molecules using the TrueSeq® Stranded mRNA library preparation kit (Illumina). The stranded mRNA library was then sequenced at a depth of 40 million reads per sample, with a total read length of 150 paired-ended base pairs (bp), using a NovaSeq 6000 S4 system (Illumina). The quality of the samples was verified using FastQC software. Trim Galore! was used for trimming ends and adapters. Genome contaminants and ribosomal RNA were removed with BBSPlit and SortMeRNA software, respectively. The alignment of reads to the mouse genome (Genome Reference: GRCm38) was performed using STAR. Gene expression quantification using read counts of exonic gene regions was carried out using Salmon. UMI-tools and Picard were used for Unique Molecular Identifiers (UMI)-based deduplication and marking, respectively. Subsequent analyses were performed using either ‘count’ or ‘tpm’ (Transcript Per Million) matrix datasets. PCA and hierarchical clustering were performed using pre-filtered datasets (removing rows with no counts, or with only a single count across all samples) after rlog transformation. DESeq2 software was used to identify the Differentially Expressed Genes (DEGs, cuttoff: Padj<0.05) and fold changes between experimental conditions. This dataset was also employed to derive gene signatures (gene sets) associated with either *Hsp90aa1* or *Hsp90ab1* silencing in CAFs or NFs, that were employed in subsequent analyses. Unless stated otherwise, expression values for each gene represented in the figures or tables were z-score normalized from the ‘tpm’ matrix dataset.

### Proteomic analyses

Cells were lysed with 6M Gu-HCl, 1 mg/mL 2-CAA and 1.5 mg/mL TCEP. Cell lysates were then boiled for 5 min at 95 °C, followed by incubation with 1 µL of LyC (Waiko) at 37 °C in a shaking heat block for 2 h. After that, 1 µL of MS-Grade Trypsin (Pierce) was added per sample and incubated o/n at 37 °C for protein digestion. The following day, 5 µL of 10% TFA was added and centrifuged for 5 min at 17,000 g to recover the supernatant with digested peptides. Stage tips were prepared by placing two Empore™ C18 extraction disks into a 200 μL pipette tip. C18 disks were then activated by adding 15 µL methanol and washed with 0.1% TFA by centrifuging 1 min at 500 rcf. Samples were then centrifuged through the disks 5 min at 500 rcf and disks were washed again with 0.1% TFA prior elution of the samples using 10% ACN and 0.05% TFA onto a 96-well plate. Samples were vacuum dried and reconstituted in LC-MS-grade H_2_O, and peptide content was estimated at 280 nm absorption using a Nanodrop 2000 spectrophotometer (ThermoFisher Scientific). For LC-MS analysis of the samples, 1 µg of peptides was injected and separated on an UltiMate™ 3000 RSLCnano UHPLCsystem (ThermoFisher Scientific), using an Aurora C18 packed emitter (IonOpticks), with a gradient from 4% to 29% ACN in 90 min and a 10 min 80% ACN wash. 0.5 % acetic acid was present throughout. The UHPLC system was coupled online with an Orbitrap Fusion™ Lumos™ Tribrid™ mass spectrometer (ThermoFisher Scientific) operated in Data-Independent Acquisition (DIA) mode, acquiring a MS 350-1650 Da at 120 k resolution followed by MS/MS on 45 windows with 0.5 Da overlap (200-2000 Da) at 30 k with a Normalized Collision Energy (NCE) setting of 27.

#### Proteomic data analysis

The raw spectra files were searched using DIA-NN software (version 1.8.1, Demichev, Ralser and Lilley labs) against the UP000000589 *Mus musculus* database, using the default setting for library-free search. Protein and peptides expression ratios were obtained and normalized using Perseus software (version 2.0.11, MaxQuant), as well as performance of the corresponding statistical t-tests for comparison between conditions. For the analysis of proteins in CDMs, a similar approach was performed, and intracellular proteins were removed from the analysis. Unless stated otherwise, expression values for each protein represented in the figures or tables were z-score normalized from the normalized data.

### Gene set enrichment analyses (GSEA)

For transcriptomic analyses by GSEA, gene count RNAseq data was first pre-processed using the ‘Voom Normalization’ from the GenePattern platform, following the program guidelines (expression value filter threshold=1). Pre-processed data was then run through GSEA software (Broad Institute of MIT and Harvard, USA) following the program guidelines. The specific settings applied in all analyses were: Number of Permutations (1000); Permutation Type (Gene set); Enrichment statistic (Weighted); Metric for ranking genes: (*t* Test). Analysed gene sets were retrieved from the Molecular Signatures Database (MSigDB). In addition, in-house manually curated gene sets or gene sets from specific publications were employed and are available upon request. Results were represented using an in-house adapted script from GSEA Multi-Sample Running Enrichment plots. Similar approaches were employed to analyse proteomics datasets using normalized proteomic data, using the gene name for the analysis. Values represent the False Discovery Rate (FDR) and the Nominal Enrichment Score (NES) of each gene set.

To calculate the gene signature score in each individual sample, we used single sample Gene Set Enrichment Analysis (ssGSEA) Projection Software from the GenePattern platform, following the programs guidelines. Each ssGSEA enrichment score represents the degree to which the genes in a particular gene set are co-ordinately up- or downregulated within a sample. ssGSEA scores were z-score normalised.

Similar approaches (GSEA and ssGSEA) were employed to investigate the enrichment and/or expression of particular gene signatures in stromal datasets.

### Gene expression analyses of clinical datasets

Gene expression analyses of human tumour stroma were retrieved from NCBI Gene Expression Omnibus (GEO). Datasets include: Finak (GSE9014, Breast), Yeoung (GSE40595, Ovary), and Nishida (GSE35602, Colon). These datasets were employed to identify consistent DEGs in cancer stroma vs normal stroma (cut-off Padj<0.05) and common up- and downregulated genes, that were employed in subsequent analyses. Additional datasets include two independent human datasets of BC stroma suitable for ssGSEA analyses: GSE8977 and GSE26910. All these datasets were also employed in other bioinformatics approaches including DESeq2, GSEA and ssGSEA, following methodology explained before. ssGSEA analysis of GSE8977, and GSE26910 was employed to investigate the Person correlation coefficient between gene signatures of interest or the normalized expression of gene signatures in each individual sample. For representation purposes, ssGSEA and gene expression values were z-score normalized. GSEA was employed in GSE9014, GSE40595 and GSE35602 to confirm the enrichment of HSP90i signatures. GSE20086 (human breast NF and CAFs), GSE70468 (human colon NF and CAFs), GSE35250 (human ovarian NF and CAFs) and GSE45256 (murine breast NF and CAFs) were also employed to assess the enrichment of HSP90i signatures in CAFs by GSEA. Finally, a HSP90i gene signature in murine embryonic fibroblasts (MEFs) was generated by identifying the top 500 downregulated DEGs after incubation with 200nM of NVP-AUY922 (HSP90 inhibitor) for 16 h (GSE125161). For microarray-based datasets, probe-to-gene collapsing was performed using the highest variance probe.

### Single cell RNAseq (scRNAseq) processing and analysis

Breast scRNAseq data was obtained from the European Genome-Phenome Archive, reference E-MTAB-10607^23^. This dataset consists of 14 fresh human breast tumour tissue samples. Each sample was pre-processed individually following the methodology of their original publication, with some changes in software versions in R. Once all individual cells were assigned to specific clusters/cell types, we assigned the expression of normalized gene counts to each cell. Individual cell gene set expression was calculated with Seurat’s AddModuleScore function. Either target gene or gene signature were projected on the Uniform Manifold Approximation and Projection (UMAP) graphs to obtain a visualization of gene expression per cluster. In addition, matrices of gene/gene signature expression per cell were further processed for additional analyses. Briefly, for correlation analyses, the Person coefficient between the expression of particular genes/gene signatures was calculated for each identified cell type. For analyses of expression in each cell cluster, the average expression per cell type was calculated and then scaled within a 1 to 0 range (1 high expression, 0 no expression). For genes, where the lowest value was always 0, average values were scaled by dividing by the highest average value. For gene signatures, where certain cell types might contain negative values, average values were scales by subtracting the minimum average value and dividing by the total range (maximum average value minus minimum average value).

### Unbiased chemical genomics approach

To identify drug treatments that could potentially regulate the expression of DEGs consistently upregulated in cancer stroma, we interrogated the gene list through the LINCS database using the “Enrichr” platform. The LINCS library has gene-expression profiles induced by over 20,000 compounds, shRNAs, and kinase inhibitors in the L1000 platform, and we specifically interrogated the “LINCS_L1000_Chem_Pert_down” database (*i.e.* drug treatments that reduce the expression of our genes of interest), using the “combined score” metric rank in the results.

### Enrichment analysis

DEGs and other gene signatures obtained from previous differential analyses were also processed for enrichment analyses using online platforms such as “DAVID” and “g:Profiler”. Briefly, “DAVID” was employed to perform functional enrichment analysis (over-representation analysis) of candidate gene lists over Gene Ontology datasets such as KEGG, GOTERM_BP and GOTERM_MF. “g:Profiler” was employed to perform functional enrichment analysis of candidate gene list over manually curated-user provided datasets, provided as “.gmt” files. Adjusted p values (-log_10_) were employed for representation.

### Integrative analysis of proteomics and transcriptomics datasets

For the integrative analysis of proteomics and transcriptomic datasets to identify HSP90α effector proteins that may be modulating YAP activity, we assumed that the potential regulator had to be directly or indirectly linked to YAP, and had to be modulated by HSP90. Thus, candidates for interrogation were initially selected based on two parameters: (i) they had to be related to YAP/TAZ regulation; and (ii) they had to present evidence of HSP90α interaction. We first generated a gene list of HSP90α interactors and YAP-related proteins. The list of HSP90α interactors was obtained from the “BioGRID” database on September 2023, obtaining 1,156 interactors. For the list of YAP-related proteins, we first obtained a similar list from BioGRID to identify all proteins reported to interact with YAP (766 proteins). This list was expanded by including all known factors associated with YAP by adding genes contained in the following gene signatures obtained from MSigDB: “GOBP HIPPO SIGNALING”, “GOBP NEGATIVE REGULATION OF HIPPO SIGNALING”, “GOBP POSITIVE REGULATION OF HIPPO SIGNALING”, “GOBP REGULATION OF HIPPO SIGNALING”, “REACTOME SIGNALING BY HIPPO”, “WP HIPPO SIGNALING REGULATION PATHWAYS”, “WP LEUKOCYTEINTRINSIC HIPPO PATHWAY FUNCTIONS”, “WP MECHANOREGULATION AND PATHOLOGY OF YAPTAZ VIA HIPPO AND NONHIPPO MECHANISMS” (917 proteins). Finally, we obtained the intersection between both lists (HSP90α interactors and YAP-related proteins) to obtain a list of 233 common elements, of which 147 were represented in the proteomics dataset and were subjected to further analysis. The proteomics expression data of those proteins was first averaged between the different replicates and z-score normalized. In addition, the ssGSEA score for the signature “NON_PROLIF_CAF_YAPTAZ_ALL_UP” obtained from the transcriptomics data for each sample was also averaged between the different replicates and z-score normalized. Finally, the Person correlation coefficient between the z-score normalized expression of each protein and the z-score normalized ssGSEA score from all samples was calculated. For the analysis with TAZ signatures, the signature “ZHANG_TAZ_GENES_NOT_YAP” was assessed in the transcriptomics data as the “NON_PROLIF_CAF_YAPTAZ_ALL_UP” signature. This signature contains genes induced by TAZ but not YAP in MCF10A cells. Candidate proteins were projected in an interaction network using STRING (https://string-db.org/).

### Survival analysis

Analysis of clinical relevance of specific genes or gene set expression was assessed using publicly available data from the Kaplan–Meir Plotter platform for breast, colon and ovarian cancer (www.kmplot.com). Probe-to-gene mapping was performed using Jetset. For survival analysis of ER negative breast (recurrence-free survival) and colon (disease-specific survival) cancer datasets, the highest tercile of gene expression was used to dichotomise the different tumours into high and low groups; for ovarian cancer (progression-free survival), “autoselect best cut-off” was selected.

In addition, we employed the METABRIC dataset to analyse the prognostic value of certain gene sets in BC patients. The gene expression data and associated clinical information were retrieved from cBioportal. Non-classified molecular subtypes, normal samples and samples with missing tumour size, grade, ER, HER2, or PR status were removed from the analysis. In all cases, survival was censored at 120 months. Patients were stratified based on median expression, and overall survival and recurrence-free survival analysed (log-rank test). Survival curves were estimated using the Kaplan-Meier method using GraphPad Prism software (GraphPad Software, Inc.). To calculate the gene signature values in each individual sample, we used ssGSEA. For this analysis, genes with no values (N/A) in any of the analysed patient samples were removed from the analysis.

### Statistical analyses

Unless specified otherwise, all statistical analysis, graphs and heatmaps were performed and generated using GraphPad Prism software (GraphPad Software, Inc.). When *n* permitted, values were tested for Gaussian distribution using the D’Agostino-Pearson normality test. For Gaussian distributions, paired or unpaired two-tailed Student’s t-test and one-way ANOVA with Tukey post-test (for multiple comparisons) were performed. For non-Gaussian distributions, Mann-Whitney’s test and Kruskal-Wallis test with Dunn’s post-test (for multiple comparisons) were performed. Unless stated otherwise, mean values and standard errors (SEM) are shown. Survival curves were estimated based on the Kaplan–Meier method and compared using a log-rank test. *P* values of less than 0.05 were considered statistically significant. Where indicated, individual *p* values are shown. Alternatively, the following symbols were used to describe statistical significance: *, *P* < 0.05; **, *P* < 0.01; ***, *P* < 0.001; #, *P* < 0.0001; n.s., non-significant.

## ACKNOWLEDGEMENTS

We thank Erik Sahai for providing us with plasmids; Maria M. Caffarel and Emad Moeendarbary for training on orthotopic mammary injections and vascularization microfluidic devices, respectively; Erik Sahai, Kristian Pietras, Hector Peinado and Daniela Quail for providing cancer cell lines; and members of the Animal Experimentation and Light Microscopy Units at IBBTEC, and the Institute of Genetics and Cancer Mass Spectrometry Facility for assistance. We also thank lab members for help and advice throughout this work. This work was funded by Spanish Government MICIU/AEI/10.13039/501100011033 (RYC-2016-20352, RTI2018-096778-A-I00, PID2021-128107OB-I00) and Asociación Española Contra el Cáncer-AECC (LABAE19044CALV). F.C. was also funded by AECC (PRYCO211372RODR); BBVA Leonardo Awards (IN[19]_BBM_BAS_0076); and the European Research Council-ERC Consolidator Grant ‘antiCAFing’ (Ref. 101045756). S.D.G. has been a recipient of a AECC PhD Studentship (PRDCA19002DOMÍ) and received an EACR Travel Grant (Ref. 791). D.C. has been a recipient of a Boehringer Ingelheim Fonds PhD Travel Grant. A.V.V. was funded by Spanish Government MICIU/AEI/10.13039/501100011033 (PID2021-125702OB-I00). The mass spectrometry equipment was funded by the MRC (MR/X01293X/1) and Wellcome Trust Multiuser Equipment Grant (208402/Z/17/Z).

## CONTRIBUTIONS

F.C. conceived the study. F.C. and S.D.G. designed the experiments. S.D.G., J.R., C.L.P. and F.C. performed and analysed the experiments. S.D.G., M.J. and F.C. performed and analysed transcriptomic analyses. M.J. and F.C. performed bioinformatic analyses. A.v.K. and S.D.G. performed and analysed proteomic analyses. S.D.G. and A.M.V. performed in vivo experiments, with help from C.L.P. and P.C. D.C. performed and analysed vascularization analyses. S.D.G. and F.C. wrote the manuscript. All authors critically read the manuscript and provided intellectual input.

## DATA AVAILABILITY

Data generated during the course of this study, including RNAseq and MS-proteomics datasets of murine NF1 and CAF1 after transfection with control, *Hsp90aa1* or *Hps90ab1* RNAi is currently being processed for deposition in appropriate public repositories.

The following public datasets have been employed in this study and acquired through NCBI GEO. Human tumour stroma: Finak (GSE9014, Breast), Yeoung (GSE40595, Ovary), and Nishida (GSE35602, Colon), Karnoub (GSE8977, Breast) and Planche (GSE26910, Breast); expression profiles of murine and human NF/CAFs: GSE20086 (human breast NF and CAFs), GSE70468 (human colon NF and CAFs), GSE35250 (human ovarian NF and CAFs), GSE45256 (murine breast NF and CAFs); expression profiles of freshly isolated murine mammary NF and CAFs (GSE195864); expression profile of MEFs subjected to control and 200 nM of NVP-AUY922 (HSP90 inhibitor) for 16h (GSE125161); expression profile and associated clinical information of colorectal cancer patients (GSE17538). METABRIC gene expression data and the associated clinical information were obtained from cBioPortal. Breast scRNAseq data was obtained from the European Genome-Phenome Archive (E-MTAB-10607).

## SUPPLEMENTARY FIGURES

**Supplementary Figure 1.**
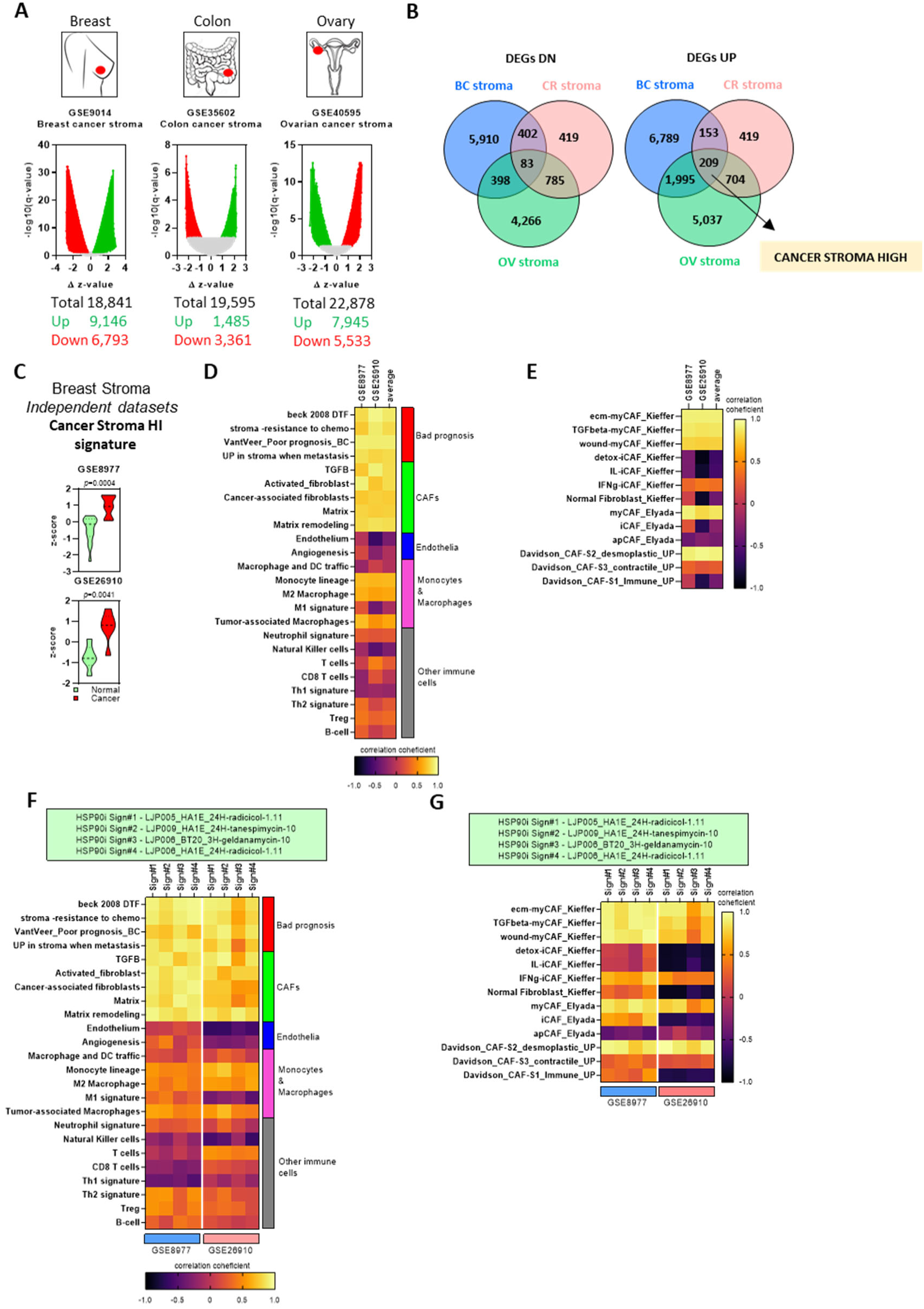
Identification of gene signatures associated with tumour stroma and CAFs. **A.** Graphs showing DEGs resulting from the gene expression analysis in breast (GSE9014), colorectal (GSE35602) and ovarian (GSE40595) databases (tumour stroma vs normal stroma). Graphs show adjusted p-value (-log_10_ q-value) and difference in z-score between cancer and normal samples. Each point represents an individual gene. **B.** Schematic representation of consistently downregulated (left) and upregulated (right) DEGs in the stroma of breast (BC), colorectal (CR) and ovarian (OV) cancer. Overlapping areas include the number of commonly upregulated or downregulated genes. **C.** Graphs showing the z-score of the ssGSEA score for the “CANCER STROMA HIGH” signature in normal and cancer stroma samples from BC patients (GSE8977: normal, n=15; cancer, n=7. GSE26910: normal, n=6; cancer, n=6). **D.** Heatmap showing the correlation coefficient between the “CANCER STROMA HIGH” signature and signatures related to immune cells, endothelia, CAFs and bad prognosis in the stromal BC datasets GSE8977 and GSE26910, as informed by ssGSEA analysis. **E.** Heatmap showing the correlation coefficient of the “CANCER STROMA HIGH” signature and signatures related to different CAF subtypes in the stromal BC datasets GSE8977 and GSE26910, as informed by ssGSEA analysis. **F.** Heatmap showing the correlation coefficient between the HSP90i signatures and signatures related to immune cells, endothelia, CAFs and bad prognosis in the stromal BC datasets GSE8977 and GSE26910, as informed by ssGSEA analysis. **G.** Heatmap showing the correlation coefficient between the HSP90i signatures and signatures related with the different CAF subtypes in the stromal BC datasets GSE8977 and GSE26910.

**Supplementary Figure 2.**
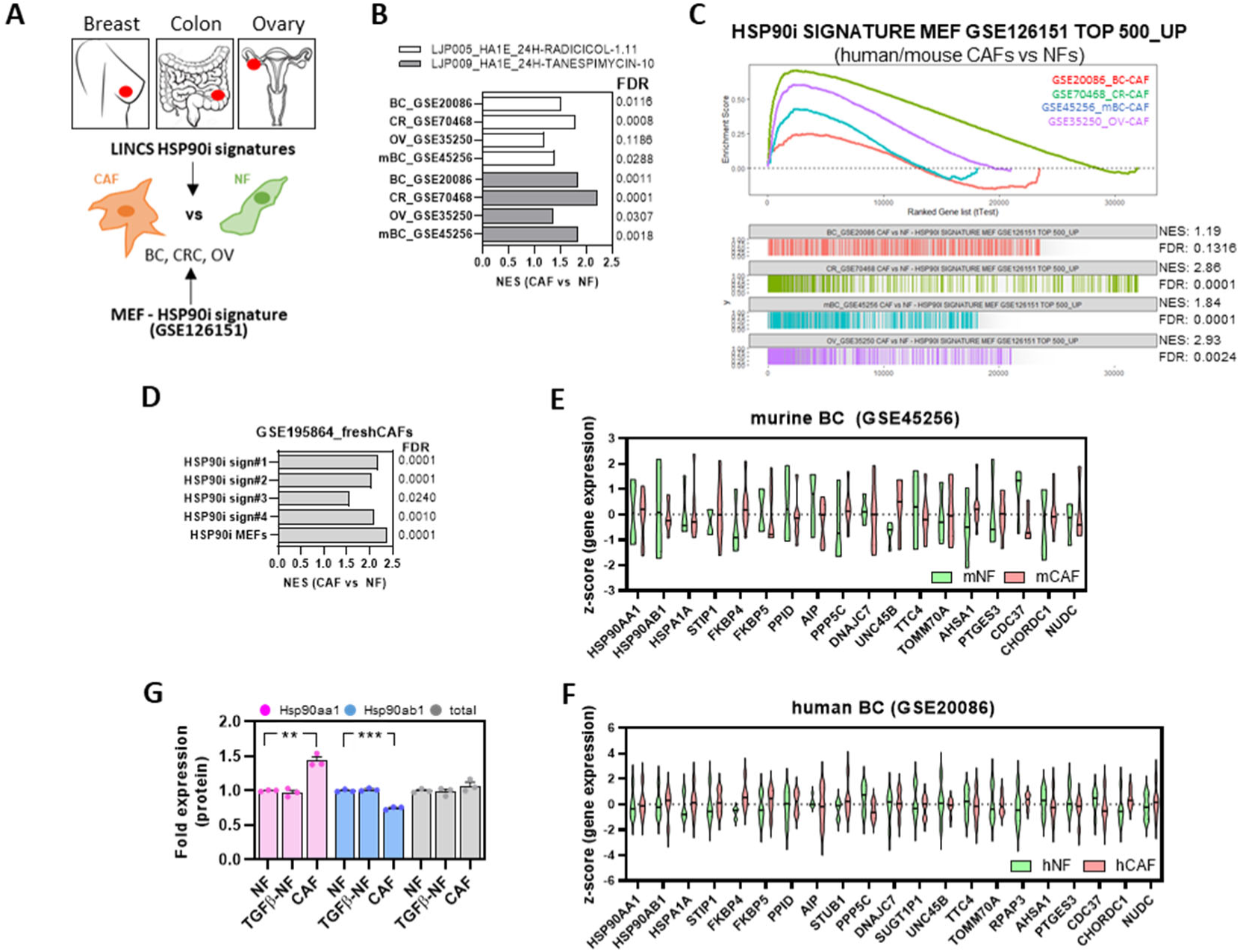
HSP90 related gene signatures are a feature of CAFs. **A.** Schematic representation of the approach to assess HSP90i-target gene enrichment in CAFs vs NF from different sources, and the methodology to generate fibroblast-specific HSP90i signatures extracted from GSE12615: mouse embryonic fibroblasts (MEFs) exposed to HSP90 inhibitor NVP-AUY922 for 16 h. **B.** Graph showing the NES and FDR scores for the enrichment of the indicated HSP90i signatures in the CAFs vs NFs of human BC (GSE20086), human colorectal cancer (GSE70468), human ovarian cancer (GSE35250) and murine BC (GSE45256), as informed by GSEA analysis. **C.** GSEA plot showing the enrichment of the indicated HSP90i signature in the different databases explored in (E). Corresponding NES and FDR values are also shown. **D.** Graph showing the NES and FDR scores for the GSEA of the different HSP90i signatures in CAFs vs NFs from the GSE195864 dataset (murine NF and CAFs freshly isolated from tissues, no in vitro expansion). **E-F.** Graphs showing the z-score gene expression of the indicated genes associated with HSP90 function in murine (E) and human (F) NFs and CAFs (Murine BC: NF, n=4; CAF, n=8. Human BC: NF, n=6; CAF, n=6). **G.** Graph showing fold protein expression levels of both HSP90 isoforms (Hsp90α and Hsp90β, respectively) and total Hsp90 in control and TGFβ1-stimulated NFs and CAFs, as informed by MS analyses (n=3). Where indicated, individual *p* values are shown; alternatively, the following symbols were used to describe statistical significance: **, *P* < 0.01; ***, *P* < 0.001.

**Supplementary Figure 3.**
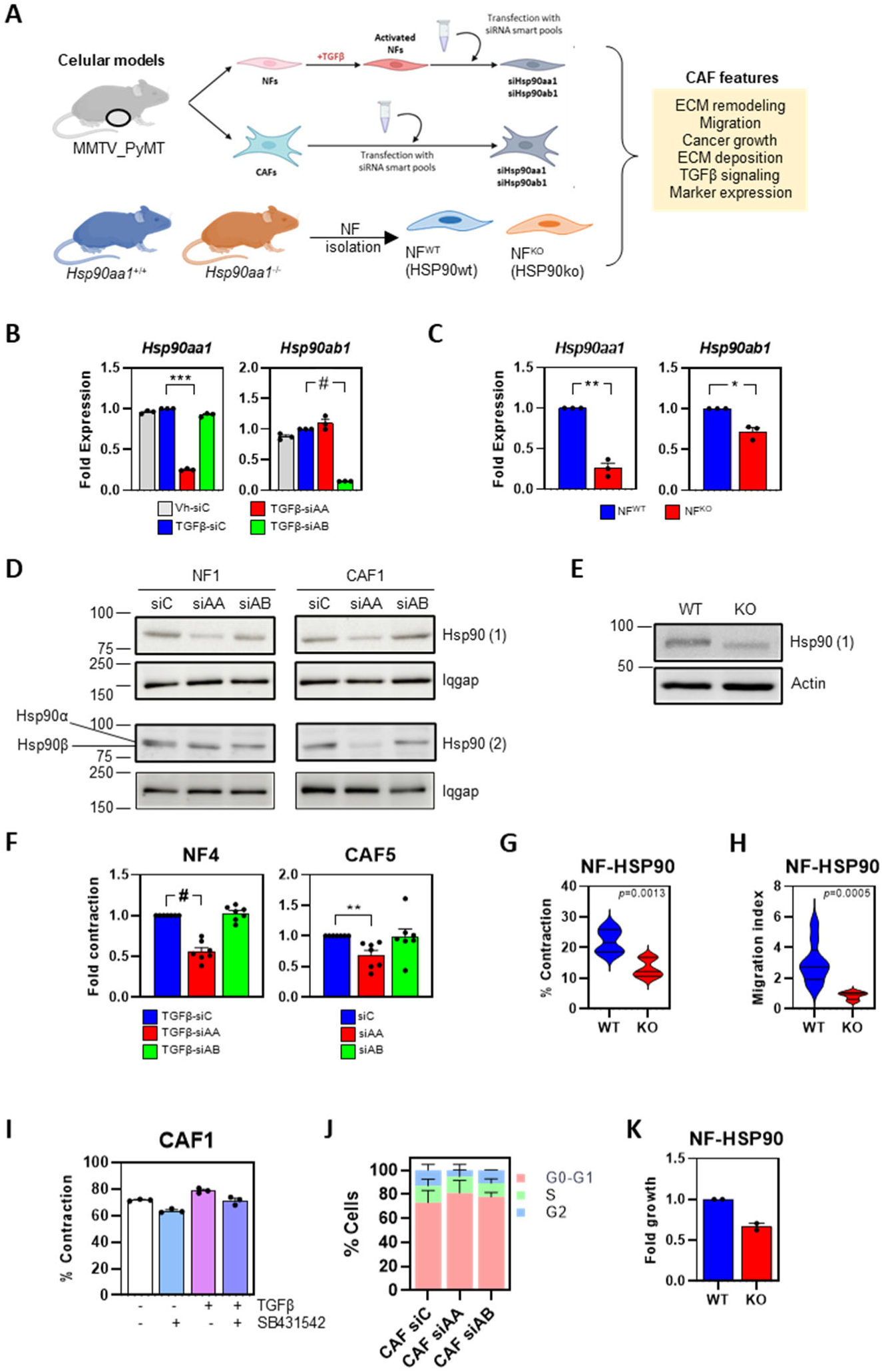
Modulation of *HSP90aa1* expression affects CAF behaviour. **A.** Schematic representation of the in vitro systems employed to assess the role of HSP90 in CAF features. All *in vitro* experiments were performed using murine CAFs and NFs transfected with the corresponding RNAi for *Hsp90aa1* and *Hsp90ab1*, or NFs derived from wild-type (NF^WT^) or *Hsp90aa1*-KO (NF^KO^) mice. **B.** Graphs show *Hsp90aa1* and *Hsp90ab1* fold mRNA expression (relative to *Gapdh*) in murine NF1 after transfection with control (siC), *Hsp90aa1* (siAA) or *Hsp90ab1* (siAB) RNAi (smart-pools). Where indicated, NF1 were stimulated with vehicle (Vh) or TGFβ1 (n=3). **C.** Graphs show *Hsp90aa1* and *Hsp90ab1* fold mRNA expression (relative to *Gapdh*) in murine NF^WT^ and NF^KO^ (n=3). **D.** Western blot showing expression levels of Hsp90 (using 2 different antibodies) and Iqgap in murine NF1 and CAF1 after transfection with control (siC), *Hsp90aa1* (siAA) or *Hsp90ab1* (siAB) RNAi (smart-pools). **E.** Western blot showing expression levels of Hsp90 (antibody#1) and Iqgap in murine NF^WT^ and NF^KO^. **F.** Graphs show fold gel contraction of TGFβ1-stimulated murine NF4 (left) and unstimulated murine CAF5 (right) after transfection with control (siC), *Hsp90aa1* (siAA) or *Hsp90ab1* (siAB) RNAi (smart-pools) (NF4: n=7. CAF5: n=7). **G.** Graph shows contraction index of unstimulated NF^WT^ and NF^KO^ (n=6). **H.** Graph shows migration index of unstimulated NF^WT^ and NF^KO^ (NF^WT^: n=9. NF^KO^: n=8). **I.** Histogram shows percentage of gel contraction of murine CAF1 subjected to vehicle (-), TGFβ1 and/or TGFβ inhibitor SB431542. **J.** Graph shows percentage of cells in each cell cycle phase for murine CAF1 cells after transfection with control (siC), *Hsp90aa1* (siAA) or *Hsp90ab1* (siAB) RNAi (smart-pools) (n=2). **K.** Graph shows the fold growth of PY8119-luc-GFP murine BC cells in co-culture with murine NF^WT^ and NF^KO^ (n=2). Where indicated, individual *p* values are shown; alternatively, the following symbols were used to describe statistical significance: *, *P* < 0.05; **, *P* < 0.01; ***, *P* < 0.001; #, *P* < 0.0001.

**Supplementary Figure 4.**
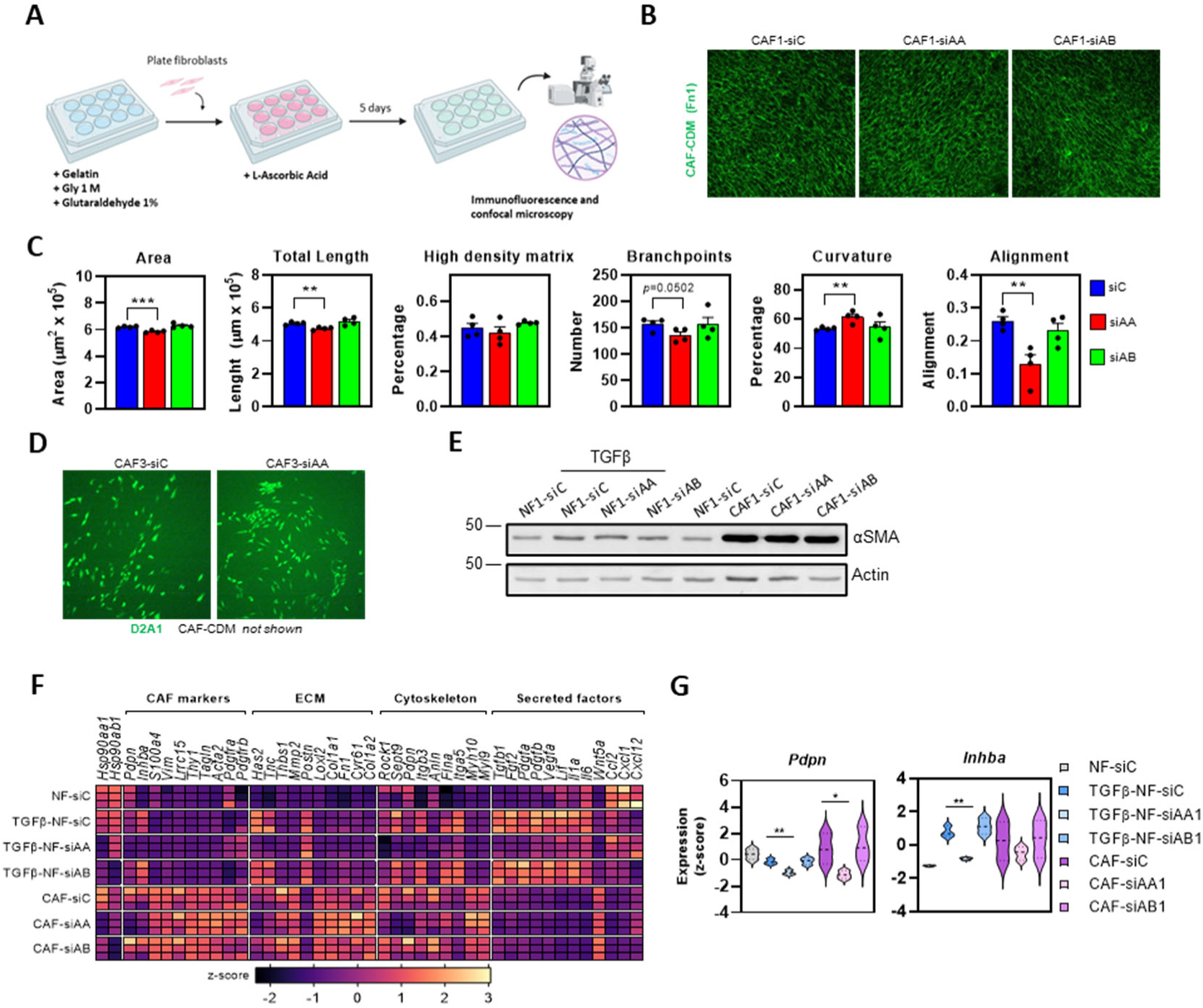
Modulation of *HSP90aa1* expression affects ECM deposition by CAFs. **A.** Schematic representation of the protocol for generating CDMs. **B.** Representative images of CDMs (Fn1, green) generated by murine CAF1 after transfection with control (siC), *Hsp90aa1* (siAA) or *Hsp90ab1* (siAB) RNAi (smart-pools). **C.** Graphs showing different parameters (area, total length of the fibres, percentage of high-density matrix, number of branch points, percentage of curvature and percentage of alignment of the fibres) from images in (B) (n=4). **D.** Representative images of D2A1-GFP cancer cells (green) cultured on top of CDMs (not shown) generated by murine CAF3 after transfection with control (siC) or *Hsp90aa1* (siAA) (smart-pools). **E.** Western blot showing expression of αSMA and Actin in murine NF1 and CAF1 after transfection with control (siC), *Hsp90aa1* (siAA) or *Hsp90ab1* (siAB) RNAi (smart-pools). Where indicated, NF1 were stimulated with vehicle (Vh) or TGFβ1. **F.** Heatmap showing the z-score expression of indicated genes, classified as CAF markers, ECM, cytoskeleton and secreted factors in murine NF1 and CAF1 after transfection with control (siC), *Hsp90aa1* (siAA) or *Hsp90ab1* (siAB) RNAi (smart-pools), as informed by RNAseq analysis in triplicates. Where indicated, NF1 were stimulated with vehicle (Vh) or TGFβ1. **G.** Graphs showing z-score of gene expression of *Pdpn* and *Inhba* in murine NF1 and CAF1 after transfection with control (siC), *Hsp90aa1* (siAA) or *Hsp90ab1* (siAB) RNAi (smart-pools). Where indicated, NF1 were stimulated with TGFβ1 (n=3). Where indicated, individual *p* values are shown; alternatively, the following symbols were used to describe statistical significance: *, *P* < 0.05; **, *P* < 0.01; ***, *P* < 0.001.

**Supplementary Figure 5.**
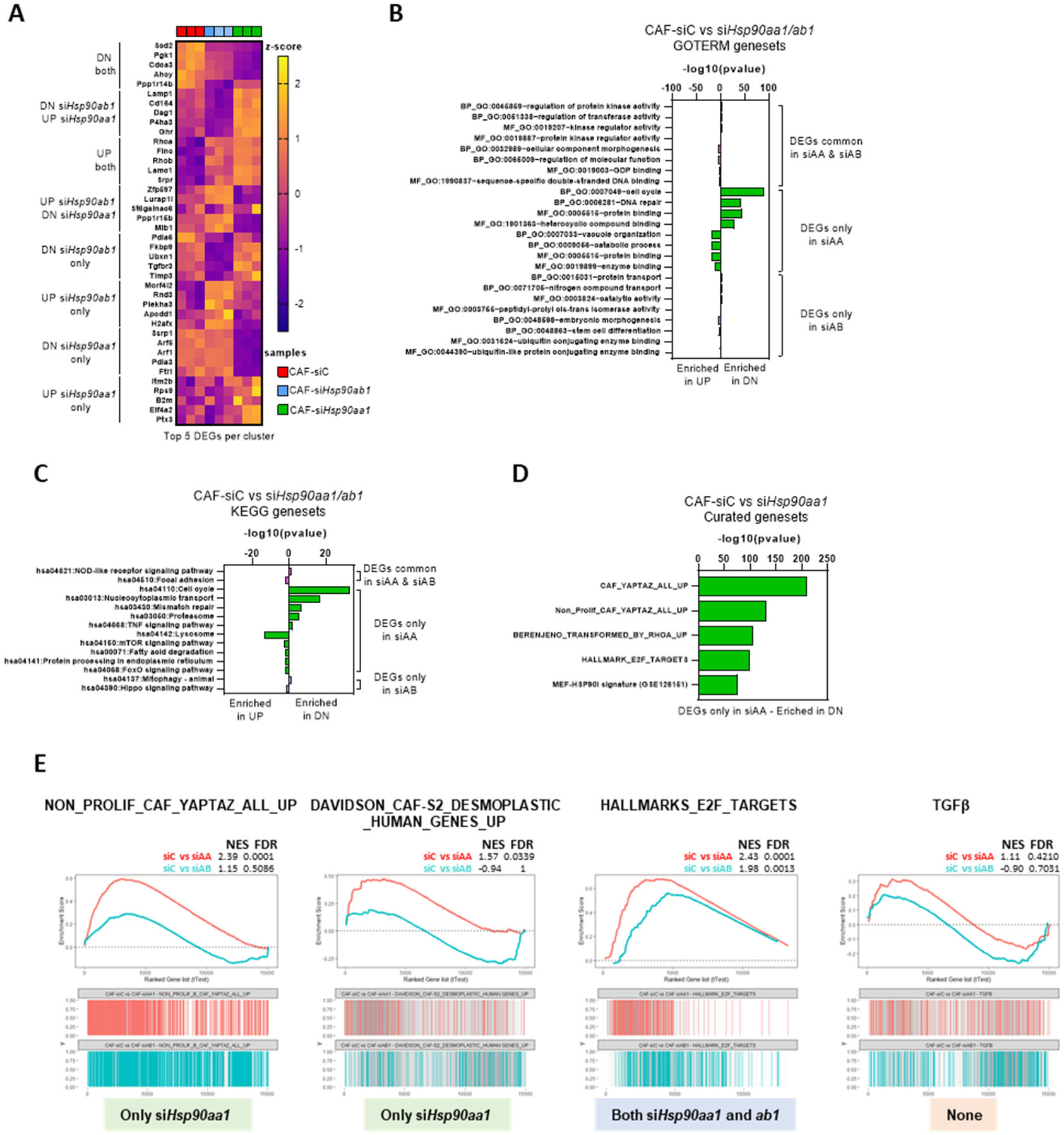
Comprehensive analysis of gene expression programs selectively modulated by HSP90α in CAF1. **A.** Heatmap showing the z-score value of the expression of the top 5 most upregulated or downregulated DEGs in CAF1 after transfection with control, *Hsp90aa1* or *Hsp90ab1* RNAi, clustered in the indicated categories (left), as informed by the RNAseq data (n=3). **B-C.** Graphs showing the enrichment score (represented as -Log10 of the P-value) in GOTERM (C) and in KEGG (D) gene sets for common and RNAi-specific DEGs found in both *Hsp90aa1* (siAA) and *Hsp90ab1* (siAB) depleted CAF1 when compared to CAF control (siC). DN represents genes significantly downregulated when compared to CAF1 control; UP represents genes significantly upregulated when compared to CAF1 control. **D.** Graph showing the enrichment score (-Log10 of the P-value) in “curated signatures” gene sets for upregulated/downregulated DEGs in *Hsp90aa1*-depleted CAF1 (siAA) when compared to CAF control (siC). **E.** GSEA plots showing the enrichment of indicated gene signatures in both *Hsp90aa1*-depleted (red line) and *Hsp90ab1*-depleted CAF1 (blue line) when compared to CAF1 control. NES and FDR coefficients are indicated. GSEAs are grouped based on their differential enrichment in the different evaluated conditions.

**Supplementary Figure 6.**
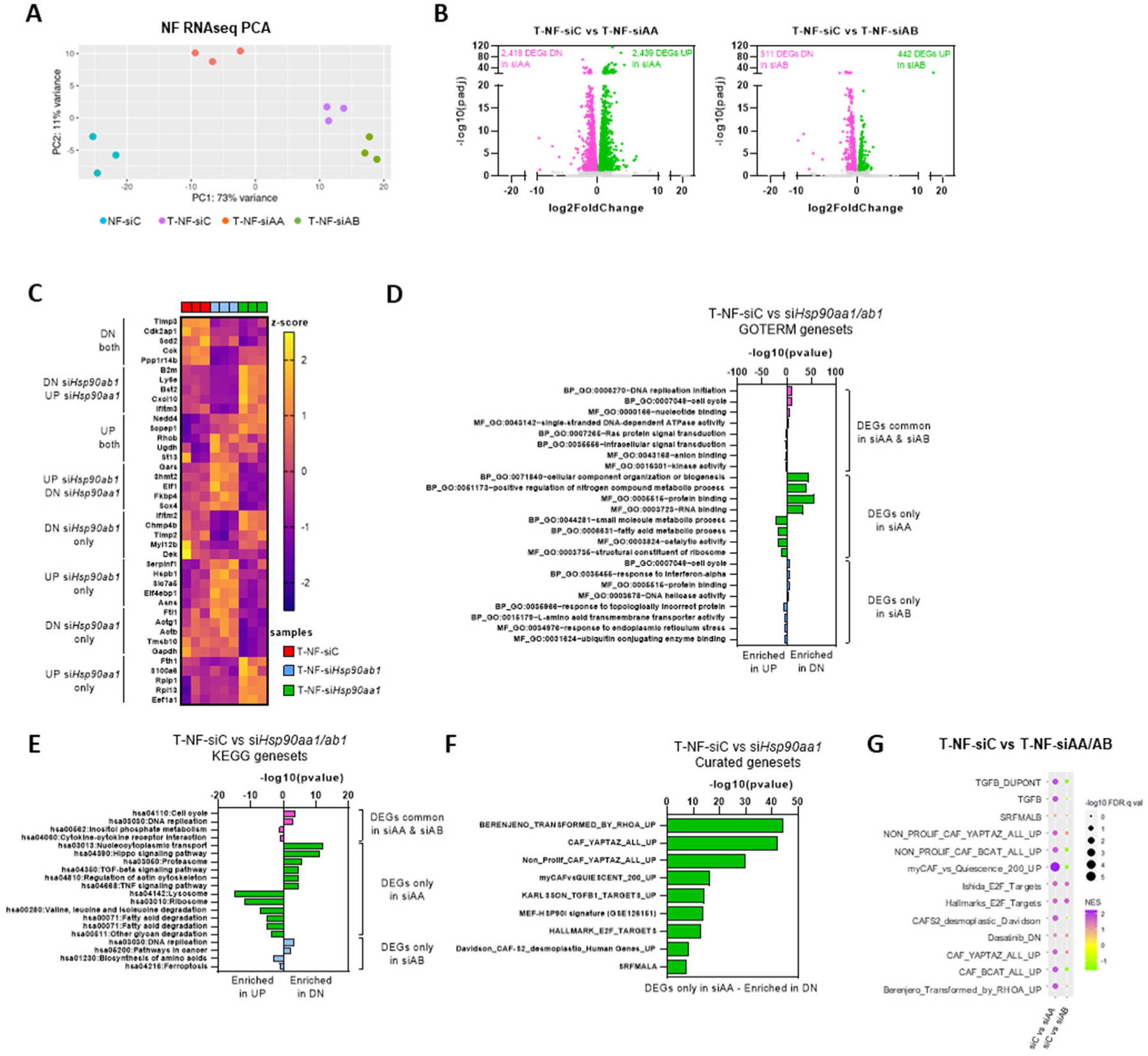
Comprehensive analysis of gene expression programs selectively modulated by HSP90α in TGFβ1-stimulated NF1. **A.** Graph showing PCA based on RNAseq of murine NF1 after transfection with control (siC), and TGFβ1-stimulated NF1 after transfection with control (T-siC), *Hsp90aa1* (T-siAA) or *Hsp90ab1* (T-siAB) RNAi (smart-pools). Each dot represents an independent replicate (n=3). **B.** Graphs showing the upregulated (green dots) and downregulated (pink dots) DEGs as determined by their -Log10 P adjusted value (Padj) and the Log2 fold change, when comparing *Hsp90aa1*-depleted (left) and *Hsp90ab1*-depleted (right) TGFβ1-stimulated NF1 (T-NF) cells, in comparison with control (siC) TGFβ1-stimulated NF1. Each dot represents an individual gene. **C.** Heatmap showing the z-score value of the expression of the top 5 most upregulated or downregulated DEGs in TGFβ1-stimulated NF1 (T-NF) after transfection with control (siC), *Hsp90aa1* or *Hsp90ab1* RNAi, clustered in the indicated categories (left), as informed by the RNAseq data (n=3). **D-E.** Graphs showing the enrichment score (represented as -Log10 of the P-value) in GOTERM (D) and in KEGG (E) gene sets for common and RNAi-specific DEGs found in both *Hsp90aa1* (siAA) and *Hsp90ab1* (siAB) depleted TGFβ1-stimulated NF1 (T-NF) when compared to TGFβ1-stimulated NF1 control (siC). DN represents genes significantly downregulated when compared to CAF1 control; UP represents genes significantly upregulated when compared to CAF1 control. **F.** Graph showing the enrichment score (-Log10 of the P-value) in “curated signatures” gene sets for upregulated/downregulated DEGs in *Hsp90aa1*-depleted TGFβ1-stimulated NF1 (siAA) when compared to TGFβ1-stimulated NF1 control (siC). **G.** Bubble plot summarizing GSEA results of “curated gene sets” when comparing TGFβ1-stimulated NF1 control (siC) vs *Hsp90aa1*-depleted (siAA) or *Hsp90ab1*-depleted (siAB) TGFβ1-stimulated NF1 cells. The size of the bubbles represents the level of significance of the signature (-Log10 FDR), and their colour represents the level of NES.

**Supplementary Figure 7.**
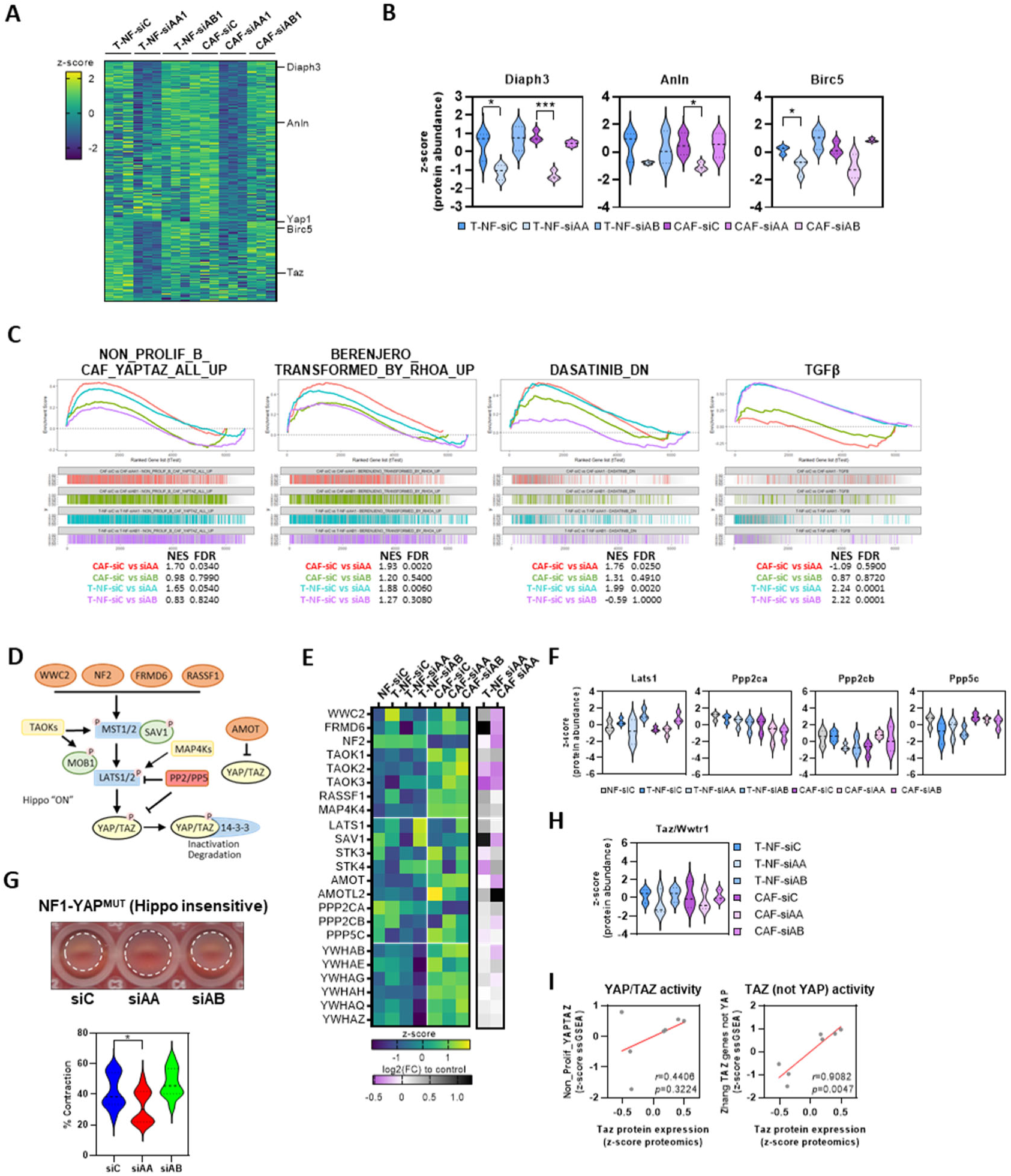
Comprehensive analysis of protein expression patterns selectively modulated by HSP90α in TGFβ1-stimulated NF1 and CAFs. **A.** Heatmap showing the z-score protein expression in CAFs and TGFβ1-stimulated NF1 (T-NF) after transfection with control (siC), *Hsp90aa1* (siAA) or *Hsp90ab1* (siAB) RNAi (smart-pools), on triplicates. Data from MS analysis. Top up- and downregulated proteins after *Hsp90aa1* silencing in both systems are shown. YAP targets are indicated. **B.** Graphs show z-score protein expression of YAP targets Diaph3, Anln and Birc5 in CAFs and TGFβ1-stimulated NF1 (T-NF) after transfection with control (siC), *Hsp90aa1* (siAA) or *Hsp90ab1* (siAB) RNAi (smart-pools)(n=3). **C.** GSEA plots showing the enrichment of the indicated signatures (based on MS-proteomics data) in *Hsp90aa1*-depleted CAF1 compared with CAF1 control (red line), *Hsp90ab1*-depleted CAF1 compared with the CAF1 control (green line), *Hsp90aa1*-depleted TGFβ1-stimulated NF1 compared with the TGFβ1-stimulated control (blue line) and *Hsp90ab1*-depleted TGFβ1-stimulated NF1 compared with the TGFβ1-stimulated control (purple line). NES and FDR coefficients are indicated. **D.** Diagram illustrating the key components of the Hippo pathway. The core components are the kinases MST1/2 (STK3/4) and LATS1/2 that promote serine phosphorylation of key residues in YAP/TAZ and in its inactivation. Phosphatases PP2/PP5 inhibit LATS1/2 phosphorylation/activation and also dephosphorylate YAP/TAZ, promoting its transcriptional activity. **E.** Heatmap showing the mean z-score protein expression value of Hippo pathway components in murine NF1 after transfection with control (NF-siC), and TGFβ1-stimulated NF1 (T-NF) or CAF1 after transfection with control (siC), *Hsp90aa1* (siAA) or *Hsp90ab1* (siAB) RNAi (smart-pools) (mean on n=3). Heatmap on the right shows the log2(FC) value of T-NF-siC vs T-NF-siAA and CAF-siC vs CAF-siAA. **F.** Graphs show z-score protein expression of selected Hippo pathway components Lats1, Ppp2ca, Ppd2cb and Ppp5c in murine NF1 after transfection with control (NF-siC), and TGFβ1-stimulated NF1 (T-NF) or CAF1 after transfection with control (siC), *Hsp90aa1* (siAA) or *Hsp90ab1* (siAB) RNAi (smart-pools) (n=3). **G.** Images and graph show gel contraction assays of NF1 cells constitutively expressing a Hippo insensitive YAP mutant (YAP-S5A) after transfection with control (siC), *Hsp90aa1* (siAA) or *Hsp90ab1* (siAB) RNAi (smart-pools)(n=12). **H.** Graph show z-score protein expression of selected Taz/Wwtr1 in murine NF1 after transfection with control (NF-siC), and TGFβ1-stimulated NF1 (T-NF) or CAF1 after transfection with control (siC), *Hsp90aa1* (siAA) or *Hsp90ab1* (siAB) RNAi (smart-pools) (n=3). **I.** Graphs show correlation between Taz expression (represented as z-score of MS-proteomics data) and the level of YAP/TAZ activity (represented as z-score of ssGSEA value of “Non_Prolif_YAPTAZ” signature from RNAseq data, left) and the level of activity of a TAZ-specific signature (represented as z-score of ssGSEA value of “Zhang_TAZ_Genes_not_YAP” from RNAseq data, right). Each dot represents the mean of 3 replicates from proteomics and RNAseq data for murine NF1 after transfection with control (NF-siC), and TGFβ1-stimulated NF1 (T-NF) or CAF1 after transfection with control (siC), *Hsp90aa1* (siAA) or *Hsp90ab1* (siAB) RNAi (smart-pools). Where indicated, individual *p* values are shown; alternatively, the following symbols were used to describe statistical significance: *, *P* < 0.05; ***, *P* < 0.001.

**Supplementary Figure 8.**
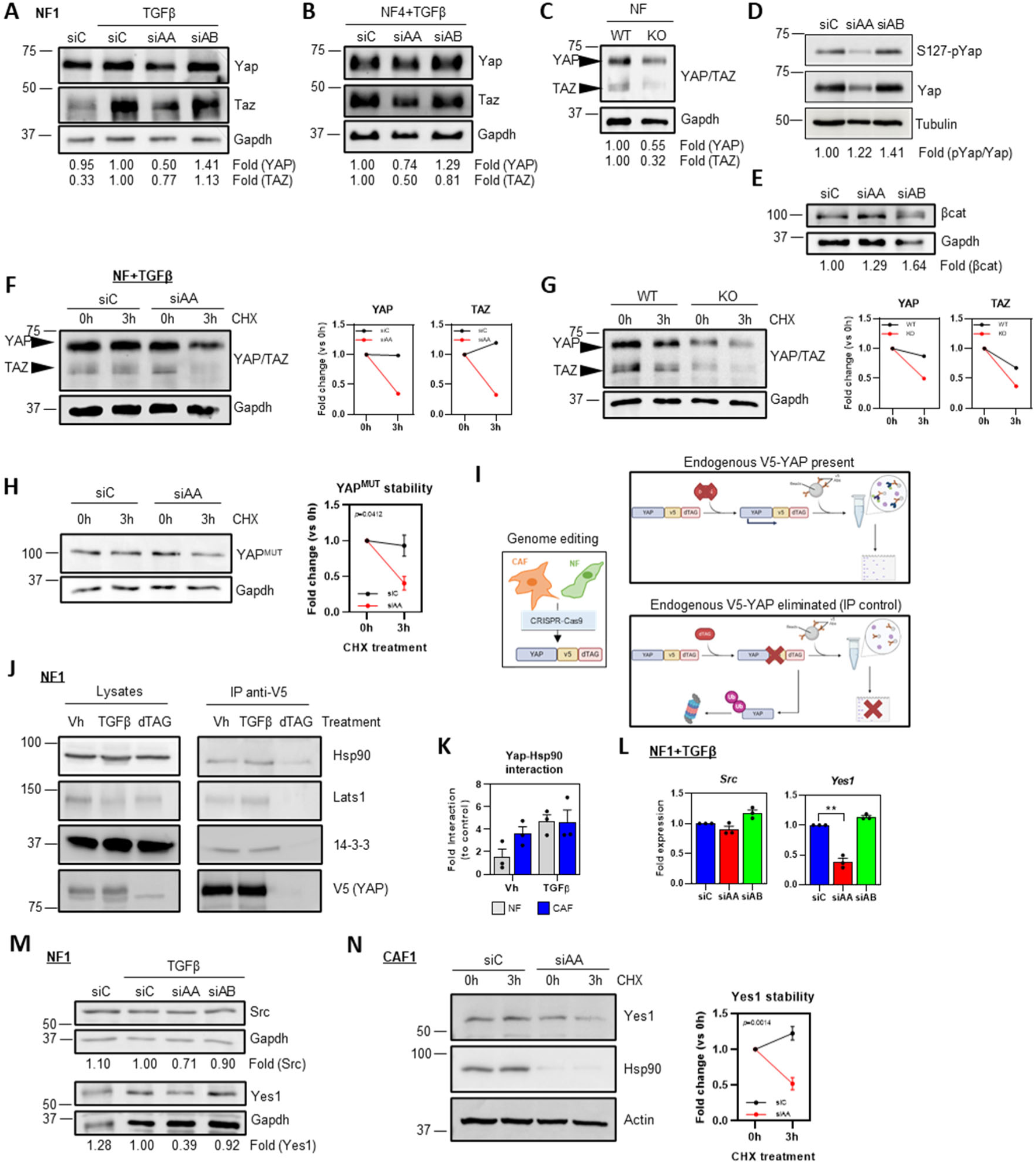
Analysis of the effect of HSP90α in YAP expression levels. **A-B.** Western blots show expression levels of Yap, Taz and Gapdh in murine NF1 (A) and NF4 (B) after transfection with control (siC), *Hsp90aa1* (siAA) or *Hsp90ab1* (siAB) RNAi (smart-pools). Where indicated, cells were stimulated TGFβ1. Fold expression levels (relative to Gapdh) are indicated. **C.** Western blots of Yap, Taz and Gapdh in unstimulated NF^WT^ and NF^KO^. Fold expression levels (relative to Gapdh) are indicated. **D-E.** Western blots show phospho-S127-Yap, Yap and Tubulin expression (D), or β-catenin (βcat) and Gapdh expression (E) in murine CAF1 after transfection with control (siC), *Hsp90aa1* (siAA) or *Hsp90ab1* (siAB) RNAi (smart-pools). Fold expression levels (relative to Tubulin or Gapdh) are indicated. **F-G.** Western blots show Yap, Taz and Gapdh expression in TGFβ1-stimulated NF1 after transfection with control (siC) or *Hsp90aa1* (siAA) RNAi (smart-pools) (F) and in the NF^WT^/NF^KO^ system (G) during pulse-chase with CHX at 0 h and 3 h. Graphs show quantification of Yap and Taz expression (relative to Gapdh) at 3 h of CHX treatment relative to the expression at 0 h for each condition. **H.** Western blot shows expression of hippo-insensitive YAP mutant (YAP^MUT^) stably expressed in murine NF1, and Gapdh after transfection with control (siC) or *Hsp90aa1* (siAA) RNAi (smart-pools) during pulse-chase with CHX at 0 h and 3 h. Graph shows quantification of YAP^MUT^ expression (relative to Gapdh) at 3 h of CHX treatment relative to the expression at 0 h for each condition (n=3). **I.** Diagram illustrating the genome editing strategy to introduce a V5-dTAG tandem in frame with the *Yap1* locus in murine NF1 and CAF1. This strategy enabled the specific immunoprecipitation of endogenous Yap with its own internal control after dTAG-induced targeted degradation. *Ub: ubiquitin.* **J.** Western blots show co-immunoprecipitation of Hsp90, Lats1 and 14-3-3 with V5-YAP (anti-V5) in NF1-V5-Yap-dTAG cells treated with vehicle (Vh), TGFβ1 or dTAG. Corresponding total lysates for all antibodies are also shown. **K.** Graph shows fold levels of interaction between Hsp90 and YAP-V5-dTAG in NF1 and CAF1 after treatment with vehicle (Vh) or TGFβ1, as inferred from co-immunoprecipitation assays in (J) and in Figure 4F. **L.** Graphs show *Src* and *Yes1* fold mRNA expression in TGFβ1-stimulated NF1 after transfection with control (siC), *Hsp90aa1* (siAA) or *Hsp90ab1* (siAB) RNAi (smart-pools). Data from RNAseq (n=3). **M.** Western blots show the expression levels of Src, Yes1 and Gapdh in murine NF1 after transfection with control (siC), *Hsp90aa1* (siAA) or *Hsp90ab1* (siAB) RNAi (smart-pools). Where indicated, cells were stimulated with TGFβ1. Fold expression levels (relative to Gapdh) are indicated. **N.** Western blot shows expression levels of Yes1, Hsp90 and Tubulin in murine CAF1 after transfection with control (siC) or *Hsp90aa1* (siAA) RNAi (smart-pools) during pulse-chase with CHX at 0 h and 3 h. Graph represents quantification of Yes1 (relative to Actin) at 3 h of CHX treatment relative to the expression at 0 h for both conditions (n=4). Where indicated, individual *p* values are shown; alternatively, the following symbols were used to describe statistical significance: **, *P* < 0.01.

**Supplementary Figure 9.**
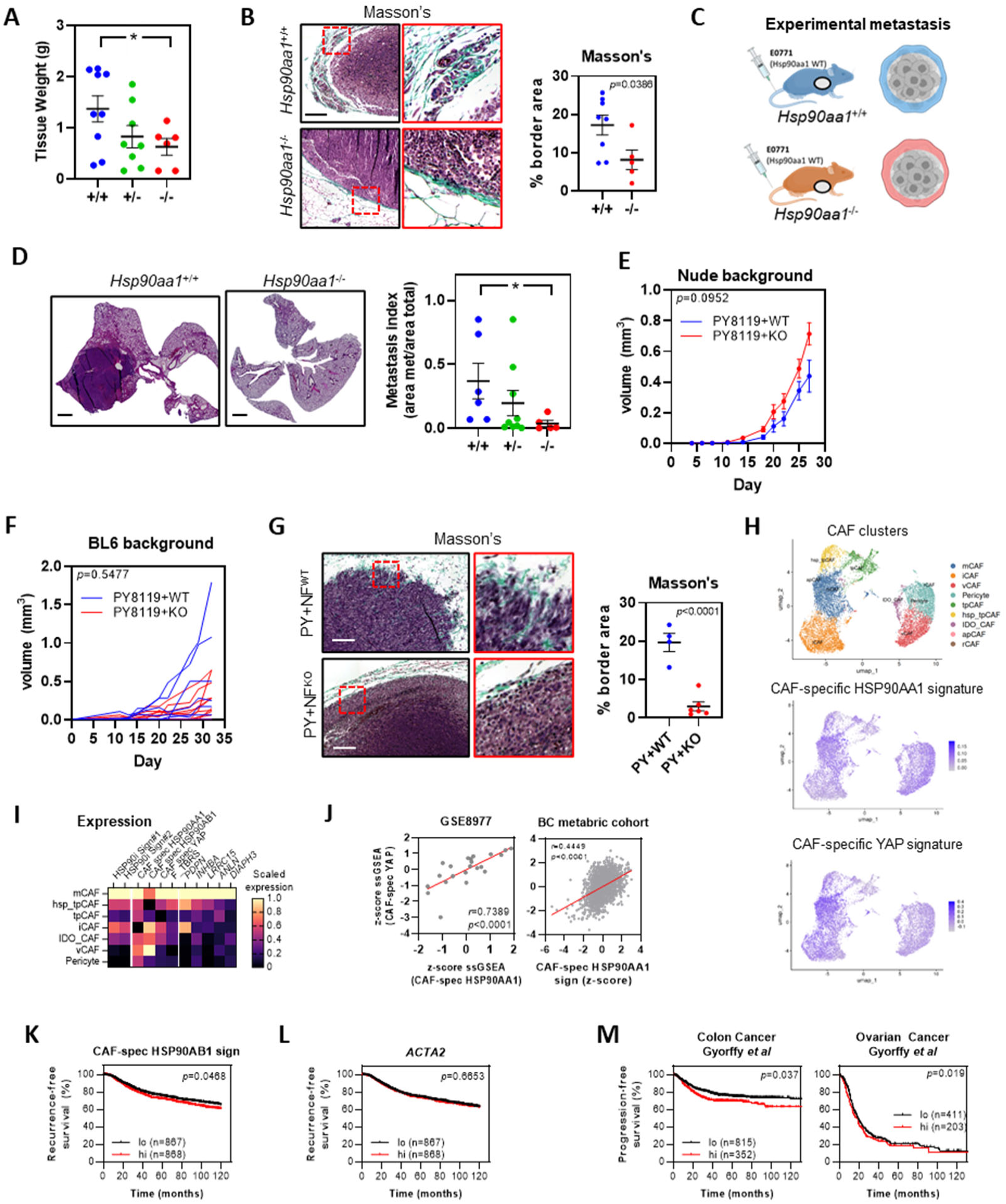
Biological role of stromal *Hsp90aa1* expression in tumour progression. **A.** Graph shows final weight of E0771 tumours growing in *Hsp90*^+/+^, *Hsp90*^+/-^ and *Hsp90*^-/-^ mice (*Hsp90*^+/+^, n=9; *Hsp90*^+/-^, n=8; *Hsp90*^-/-^, n=6). **B.** Images show Masson’s trichrome staining (representing fibrillary collagen) in sections from primary E0771 tumours (tumour border regions) grown in *Hsp90*^+/+^ and *Hsp90*^-/-^ mice. Red square represents zoom-up regions shown in right panels. Scale bar, 100 μm. Graph on the right shows the percentage of border area positive for fibrillary collagen (+/+, n=8; -/-, n=5). **C.** Schematic of the experimental metastasis model. E0771 murine BC cells (wild-type for *Hsp90*) were injected on the tail vein of *Hsp90*^+/+^, *Hsp90*^+/-^ and *Hsp90*^-/-^ adult females (BL6 background, immunocompetent) to generate lung metastases. **D.** Images show H&E staining of sections from lungs of *Hsp90*^+/+^ and *Hsp90*^-/-^ animals injected with E0771 cells. Scale bar, 1 mm. Graph on the right shows the metastasis index on lungs of *Hsp90*^+/+^, *Hsp90*^+/-^ and *Hsp90*^-/-^ animals injected with E0771 cells (*Hsp90*^+/+^, n=6; *Hsp90*^+/-^, n=9; *Hsp90*^-/-^, n=5). **E.** Graph shows tumour growth curves of subcutaneous tumours grown in nude mice after injection of metastatic MMTV-PyMT murine cancer cells (PY8116) together with NF^WT^ and NF^KO^ cells (PY+NF^WT^, n=4; PY+NF^KO^, n=6). **F.** Graph shows individual tumour growth curves of subcutaneous tumours grown in BL6 background (immunocompetent) mice after injection of metastatic syngeneic MMTV-PyMT murine cancer cells (PY8116) together with NF^WT^ and NF^KO^ cells (PY+NF^WT^, n=7; PY+NF^KO^, n=12). **G.** Images show Masson’s trichrome staining (representing fibrillary collagen) in sections from primary PY8116+NF^WT^ (PY+NF^WT^) or PY8116+NF^KO^ (PY+NF^KO^) tumours grown in nude mice. Scale bar, 100 μm. Red square represents zoom-up regions shown in right panels. Scale bar, 100 μm. Graph on the right shows the percentage of border area positive for fibrillary collagen (PY+NF^WT^, n=4; PY+NF^KO^, n=6). **H.** UMAPs showing the different CAF subtypes identified in BC patients (top), and the expression matrices for gene expression signatures for HSP90AA1 in CAFs (CAF-specific HSP90AA1 signature, center) and for YAP activity in BC CAFs (CAF-specific YAP signature, bottom), as informed by scRNAseq analysis^23^. Each dot in the graph represents a single cell. **I.** Heatmap showing scaled values of gene expression levels of the indicated gene signatures and genes in each CAF cluster type from (H). **J.** Graphs show correlation between signatures for YAP activation in CAFs (“CAF-spec YAP”), and gene expression signatures for HSP90AA1 in CAFs (“CAF-spec HSP90AA1”) in BC stroma (GSE8977) and whole tissue (METABRIC). Signatures were assessed by ssGSEA and then z-score normalized. Each point in the graph represents an individual patient. **K, L.** Graphs show percentage of recurrence-free survival of BC patients from the METABRIC dataset with high (hi) or low (lo) expression of a gene signature associated with HSP90AB1 activity in CAFs (K) or on expression of *ACTA2* (L). **M.** Graphs showing the percentage of progression-free survival patients with high (hi, red) or low (lo, black) expression of CAF-specific HSP90AA1 signature in colon (left) and ovarian cancer patients (right). Where indicated, individual *p* values are shown; alternatively, the following symbols were used to describe statistical significance: *, *P* < 0.05.

## SUPPLEMENTARY MOVIES

**Supplementary Movie 1. D2A1-GFP sphenorids over CDMs generated by CAF transfected with control RNAi, example 1.** Time-lapse movie of D2A1-GFP (green) cancer spheoid #1 recorded over 48 h when seeded over CDMs generated by CAF transfected with control RNAi (not shown).

**Supplementary Movie 2. D2A1-GFP sphenorids over CDMs generated by CAF transfected with control RNAi, example 2.** Time-lapse movie of D2A1-GFP (green) cancer spheoid #2 recorded over 48 h when seeded over CDMs generated by CAF transfected with control RNAi (not shown).

**Supplementary Movie 3. D2A1-GFP sphenorids over CDMs generated by CAF transfected with *Hsp90aa1* RNAi, example 1.** Time-lapse movie of D2A1-GFP (green) cancer spheoid #1 recorded over 48 h when seeded over CDMs generated by CAF transfected with *Hsp90aa1* RNAi smartpool (not shown).

**Supplementary Movie 4. D2A1-GFP sphenorids over CDMs generated by CAF transfected with *Hsp90aa1* RNAi, example 2.** Time-lapse movie of D2A1-GFP (green) cancer spheoid #2 recorded over 48 h when seeded over CDMs generated by CAF transfected with *Hsp90aa1* RNAi smartpool (not shown).

## SUPPLEMENTARY TABLES

**Supplementary Table 1.** Genes consistently Upregulated/Downregulated in cancer stroma (breast, colon, ovary).

**Supplementary Table 2.** Genes expression signatures.

**Supplementary Table 3.** RNAi.

**Supplementary Table 4.** Antibodies. **Supplementary Table 5.** Oligonucleotides.

